# SL/Kh Pre-B Lymphomas Originate in the Thymus and are a Model for Primary Mediastinal (Thymic) Large B-Cell Lymphoma

**DOI:** 10.1101/2024.12.16.626697

**Authors:** Tami L. Thomae, Si Hui Tan, Keiko Akagi, Jerrold M. Ward, Neal G. Copeland, Nancy A. Jenkins

## Abstract

SL/Kh mice develop a high frequency of retrovirally-induced pre-B lymphomas at 3-6 months of age. They also exhibit an abnormal transient expansion of pre-B cells in the bone marrow, although the relevance of this expansion for lymphomagenesis has remained unclear. Here, we use a dual approach that combines pathology with flow cytometry to more fully characterize the nature and origin of SL/Kh lymphomas. Unexpectedly, our studies showed that SL/Kh lymphomas arise from a rare population of pro/pre-B cells in the thymus. We also identified a 10-fold reduction in Notch1 expression in SL/Kh thymic T cells that is associated with a block in early T cell development, a reduction in the number of thymic T cells with age, and an expansion of thymic pro/pre-B cells. This phenotype is consistent with previous studies showing that Notch1 signaling is essential for lymphoid progenitors to undergo T cell commitment and for suppressing B cell development in the thymus. We propose that this developmental defect provides a niche for early B cells to accumulate in the thymus, which, when combined with subsequent retroviral insertional mutagenesis, results in the induction of pre-B lymphomas that originate in the thymus. This is also consistent with our analysis of the genes insertionally mutated in SL/Kh lymphomas, which shows that many function in signaling pathways such as JAK/STAT and RAS/MAPK/ERK that are commonly deregulated in B-cell lymphomas. Primary human mediastinal large B-cell lymphoma (MLBCL) is another lymphoma that is derived from thymic B cells, although virtually nothing is known about the cause of this rare disease. Our studies provide new insights into an underappreciated class of B-cell lymphomas and a mouse model for the study of MLBCL.

## Introduction

Inbred mouse strains were developed in part to determine if cancer was genetically determined. Scientists thought that if inbred strains with a high spontaneous incidence of cancer could be created though selective inbreeding, this would prove that cancer was genetically determined. In 1921, Carleton E. MacDowell created an inbred strain, C58, which had a high spontaneous incidence of T-cell lymphoma^1^. Shortly thereafter, Jacob Furth created the AKR inbred strain, which also had a high spontaneous incidence of T-cell lymphoma^2^. Many scientists hailed these findings because they believed this proved the genetic origin of cancer. Unfortunately, these claims turned out to be wrong.

Through the work of many different laboratories it was later shown that an infectious ecotropic murine leukemia virus (MuLV) encoded in germline of C58 and AKR mice was responsible for inducing lymphomas in these mice through a process referred to as insertional mutagenesis^3–8^. Insertional mutagenesis occurs as part of the normal life cycle of the retrovirus when the proviral DNA integrates into the host genome. During the process of integration the provirus by chance can integrate upstream or in the 5’ end of a proto-oncogene and deregulate its expression, and thereby lead to cancer. Alternatively, the provirus can integrate within the coding region of a tumor suppressor gene and inactivate its normal expression and function, and thereby lead to cancer. The value of insertional mutagenesis as a disease causing mechanism is that the cancer genes in tumors are tagged by the presence of the provirus and can thus be readily identified. During the last two decades hundreds of cancer genes have been identified by insertional mutagenesis making this one of the most powerful cancer gene discovery tools yet discovered.

Unfortunately, for reasons still not completely understood, most of the tumor prone strains created in the early years by mouse geneticists developed T-cell lymphomas. This made it impossible to use insertional mutagenesis to identify genes involved in other types of hematopoietic cancer, such as B-cell lymphoma and myeloid leukemia. In 1982, Hiai and colleagues reported the creation of a new lymphoma prone strain, SL/Kh, which was derived from the F_2_ cross of two pre-existing lymphoma prone strains, SL and AKR^9^. SL/Kh mice were different, however, because they did not die from T cell lymphomas. Two types of lymphoma were observed in SL/Kh mice: the major type (84%) involved primarily the lymph nodes and the spleen, but also often the liver^10, 11^. The minor type (16%) involved primarily the bone marrow^10, 11^. The surface phenotype of both lymphomas was indistinguishable: they expressed B220, 6C3, and c-kit but not Thy1.1, Mac-1 or surface Ig. The immunoglobulin heavy chain was also clonally rearranged but the lambda light chain was in the germ line configuration. In addition, RT-PCR demonstrated that lambda-5, RAG-1 and RAG-2 were expressed, which are specifically associated with pre-B lymphocytes. These observations indicated that SL/Kh lymphomas were pre-B-lymphomas^11, 12^.

Subsequent genetic studies showed that SL/Kh mice are susceptible to pre-B-lymphomas because of a unique combination of genes they inherited from their parents. For example, SL/Kh mice inherited the endogenous ecotropic murine leukemia provirus, *Emv11*, from the AKR parent, which is the source of the infectious MuLV responsible for insertional mutagenesis and lymphoma induction in SL/Kh mice. SL/Kh mice also carry a recessive lymphoma latency acceleration (*lla*) allele that maps to the major histocompatability locus, which accelerates the development of lymphomas induced by *Emv11* ^13^. Finally, SL/Kh mice failed to inherit the dominant thymic lymphoma susceptible mouse-1 (*Tlsm1*) allele carried by AKR, which predisposes AKR mice to T-cell disease^14, 15^.

An abnormal transient expansion of pre-B cells has also been documented in the bone marrow of SL/Kh mice, which reaches a peak at 4-6 weeks of age^12^. Such an expansion is not normally found in other strains of mice, but is observed in mice with conditions predisposing them to B-cell lymphocyte lymphomagenesis, such as Eμ-myc transgenic mice and Abelson virus-injected mice^16, 17^. Subsequent reciprocal chimeras involving BALB/c and SL/Kh mice showed that this pre-B-cell expansion is a property of the SL/Kh hematopoietic stem cells rather than the bone marrow microenvironment. Pre-B-cell expansion was also observed in F_1_ hybrid mice carrying a dominant *Fv-4^r^*retroviral restriction locus and was not inhibited by neonatal injection of maternal resistance factor which blocks retroviral spread, indicating that pre-B-cell expansion is not dependent on virus replication^18^. Genetic mapping studies subsequently identified a highly significant quantitative trait locus named bone marrow pre-B-1 (*Bomb1*) that maps to mouse chromosome 3, which is primarily responsible for this pre-B-cell expansion^19^.

Pre-B cell expansion *per se* is not, however, sufficient for SL/Kh lymphomagenesis since NFS.SL/Kh-*Bomb1* congenic mice, which also show the pre-B cell expansion phenotype, do not develop pre-B lymphomas, even after 1 year of observation, irrespective of MuLV inoculation^19^.

Despite all of the elegant genetic studies that have been conducted on SL/Kh mice over the years it is still not clear where the pre-B lymphomas observed in SL/Kh mice originate in the bone marrow, lymph nodes, spleen or elsewhere. The relevance of the early pre-B cell expansion observed in SL/Kh mice to lymphomagenesis also remains unclear, although genetic studies suggest that pre-B cell expansion *per se* is not critical for lymphomagenesis.

In studies reported here we provide data indicating that SL/Kh lymphomas arise from a rare population of B cells located in the thymus and are thus a mouse model for a rare human B-cell lymphoma that also arises from B cells located in the thymus called mediastinal large (thymic) B-cell lymphoma. We also identify a previously unrecognized defect in T-cell development in SL/Kh mice that is associated with reduced expression of Notch1, a key lineage-determining gene that promotes development of the T-cell lineage at the expense of the B cell lineage, and show how this development defect could explain the unusual thymic origin of SL/Kh pre-B cell lymphomas as well as the pre-B-cell expansion that occurs in the bone marrow of young SL/Kh mice.

## Materials and Methods

### Source of the Mice

SL/Kh mice were obtained as a gift from Dr. H. Hiai, Kyoto University, Kyoto, Japan, in early 2000, and then maintained for several years in our breeding colony at the National Cancer Institute, Frederick, Maryland. This breeding colony was subsequently transferred to the Institute of Molecular & Cell Biology (IMCB), Singapore, in 2006, where it has been maintained ever since. C57BL6/J mice were purchased from The Jackson Laboratory, Bar Harbor, Maine, and are also being maintained in our breeding colony at the IMCB.

### Histology and Pathology

Full necropsies were performed on all SL/Kh and C57BL6/J mice, with particular attention paid to the primary and secondary lymphoid organs. Necropsy and histopathology were performed on day 11-15, 25-30, 45-50, 60, 90, 120 and 180+ old (moribund) mice. At least 3 males and 3 females from each strain at ages 11 days to 120 days were used for analysis. Forty mice were allowed to develop clinical signs of disease and become moribund. Tissues were harvested and fixed in 10% neutral buffered formalin, embedded in paraffin and sectioned. Five µM sections were stained with hematoxylin and eosin or the antibody indicated. The antibodies used were Tdt (AbD Serotec #AHP59HT, Dilution 1:50), Pax5 (Novocastra #NCL-L-PAX5, Dilution 1:100), CD45R (BD Pharmingen #553086, Dilution 1:500) and CD3 (Dako # A0452, Dilution 1:500). TUNEL assays were performed using the ApopTag^®^ Plus Peroxidase Kit (Chemicon #S7101) with hematoxylin as a counter-stain.

### Tissue Dissociation

Spleen and thymus were harvested from at least three independent SL/Kh or C57BL6/J mice at 45, 60, 90, and 120 days of age. Tissues were subsequently processed to single cell suspensions using mechanical dissociation. Briefly, the organs were freshly harvested and placed into 10 ml of PBS + 2% FBS. Two slides wet with ice cold medium were used to gently disrupt the tissues on ice. For lymphocyte isolations from the femur, bone marrow plugs were flushed with 1X PBS using a 25-gauge needle. The cell suspensions were passed over a 40 µm nylon strainer (BD Falcon #352340). All cell suspensions were centrifuged at 1200 rpm for 10 minutes at 4°C. The cells were washed with 10 ml cold 1X PBS + 1% BSA (FACS buffer) and passed over a 40 µm nylon strainer and centrifuged to collect the cells. Lymphocytes were counted in the presence of trypan blue to exclude dead cells.

### Characterization of B and T cell Populations

Antibody cocktails were used to characterize the B and T cell populations in the thymus, spleen and bone marrow of the SL/Kh and the C57BL6/J mice. The antibody cocktails used were; Mouse T Lymphocyte Activation Antibody Cocktail (BD Pharmingen #557916), Mouse T Lymphocyte Subset Antibody Cocktail (BD Pharmingen #558431), Mouse B Lymphocyte Activation Antibody Cocktail (BD Pharmingen #558064), and Mouse B Lymphocyte Subset Antibody Cocktail (BD Pharmingen #558331). Cells were labeled according to the manufacturer’s directions and red blood cells were eliminated using lysing buffer (BD Biosciences #555899). Isotype controls were used for each labeling experiment. Cells were resuspended in 500 µl of FACS buffer and immediately used for flow cytometric analysis.

### Sorting of Thymic T Cells

To investigate the thymic T cell subpopulations of SL/Kh and C57BL6/J mice, markers for CD4 (BD Pharmingen #552051), CD8 (BD Pharmingen #553032), CD44 (BD Pharmingen #559250) and CD25 (BD Pharmingen #553071) were used. Single cell suspensions were generated as described above using 45-day-old thymus from SL/Kh and C57BL6/J mice. 10^8^ cells were used in triplicate for each mouse strain. Cells were blocked in mouse serum (Millipore #S25-10ml) for 15 minutes. Cells were then washed and incubated on ice for 40 minutes with the CD4 (50 µl), CD8 (50 µl), CD44 (25 µl) and CD25 (25 µl) antibodies. After labeling, cells were washed with FACs buffer and resuspended in 1X PBS + 0.1% propidium iodide. Cells were then immediately used in the sorting experiment. Sorted cells were collected and RNA was extracted for expression analysis.

### Analysis of Immunofluorescence Data

Data collection of the antibody cocktail-labeled T and B cell populations was performed using a BD FACSCalibur Flow Cytometer (#342975) running the BD CellQuest^TM^ Pro Software (Version 5.2.1). Data was analyzed using FlowJo Software (Version 4.0.2). Thymic T cell subpopulations were collected using a BD FACSVantage SE running the CellQuest^TM^ Software (Version 3.3). Data was analyzed using FlowJo Software (Version 4.0.2).

### Quantitative Real-Time PCR Expression Analysis

Total cellular RNA was extracted from sorted lymphocyte samples (20,000-200,000 cells/sample) or single cell suspensions (10^8^ cells) using TRIzol® Reagent (Invitrogen #15596-018). Total RNA was purified using the RNeasy MinElute Cleanup Kit (Qiagen #74204). Oligo-dT (Invitrogen #18418-020) primed cDNA samples were prepared from total RNA using Superscript III Reverse Transcriptase (Invitrogen #18080-044).

Quantitative-PCR analysis was performed using SYBR® GREEN PCR master mix (Applied Biosystems #4309155) using a 7500 RealTime PCR System (Applied Biosystems). Primers were designed with the Primer Express Software (Version 3.0) sequences available upon request.

### Splinkerette PCR Amplification, Cloning and Sequencing

Splinkerette PCR amplification was performed as previously described ^20^. Briefly, genomic DNA isolated from tumor samples of animals with lymphoma, was digested with *BfaI* (5’-end cloning) or *NlaIII,* (3’-end cloning), and ligated to the splinkerette linker overnight. The ligation reaction was digested with *BtsI* and *SpeI* or *BglII* to prevent 5’ or 3’ long terminal repeat (LTR) amplification, respectively (NEB). Nested PCR was then performed on ligation reactions, using splinkerette-specific primers and primers recognizing the LTR of AKR murine-leukemia-virus-inducing locus (*Emv11*). PCR products were cloned into the pCR4-TOPO vector (Invitrogen # K4575-40). DNA was then prepped from 96 colonies from each of the *NlaIII-* and *BfaI-*digested tumors using the Direct prep 96 miniprep kit (Qiagen #27361). Products were subsequently sequenced using the BigDye® Terminator v3.1 Cycle Sequencing Kit and ABI 3730xl DNA Analyzers (Applied Biosystems, Foster City, CA). Primer sequences are available upon request. Sequences were screened for LTR elements from sequence reads using cross-match. Subsequently, the vector and LTR elements were trimmed from the reads and aligned against the mouse reference genome assembly (mm9). The alignment output (90% identity/90% coverage) was parsed and the consolidated reads were placed into insertion clusters for each tumor. Potential endogenous insertions are filtered from the output based on exact nucleotide matches between tumors.

## Results

### SL/Kh Lymphomas arise from Thymic B Cells

In order to determine the origin of SL/Kh pre-B lymphomas we performed a comprehensive histopathological analysis of SL/Kh and control C57BL/6J (B6) hematopoietic tissues, beginning early in life and concluding at the time the mice developed frank lymphomas. Examination of the thymuses of SL/Kh and B6 mice at 11, 15, 25, 29-31, 45, 48 and 50 days of age showed that they were all grossly and histologically normal. A slight atrophy (loss of mostly cortical thymocytes) could sometimes be observed in the thymus of SL/Kh mice at 60 days of age, which, by 90 days, was more pronounced and affected both the medulla and cortex (Figure 1). At 120 days, the normal architecture of the SL/Kh thymus was completely obliterated. The 120-day-old SL/Kh thymus also appeared slightly larger than the 90-day-old SL/Kh thymus (Figure 1). There were no sex differences.

**Figure 1.**
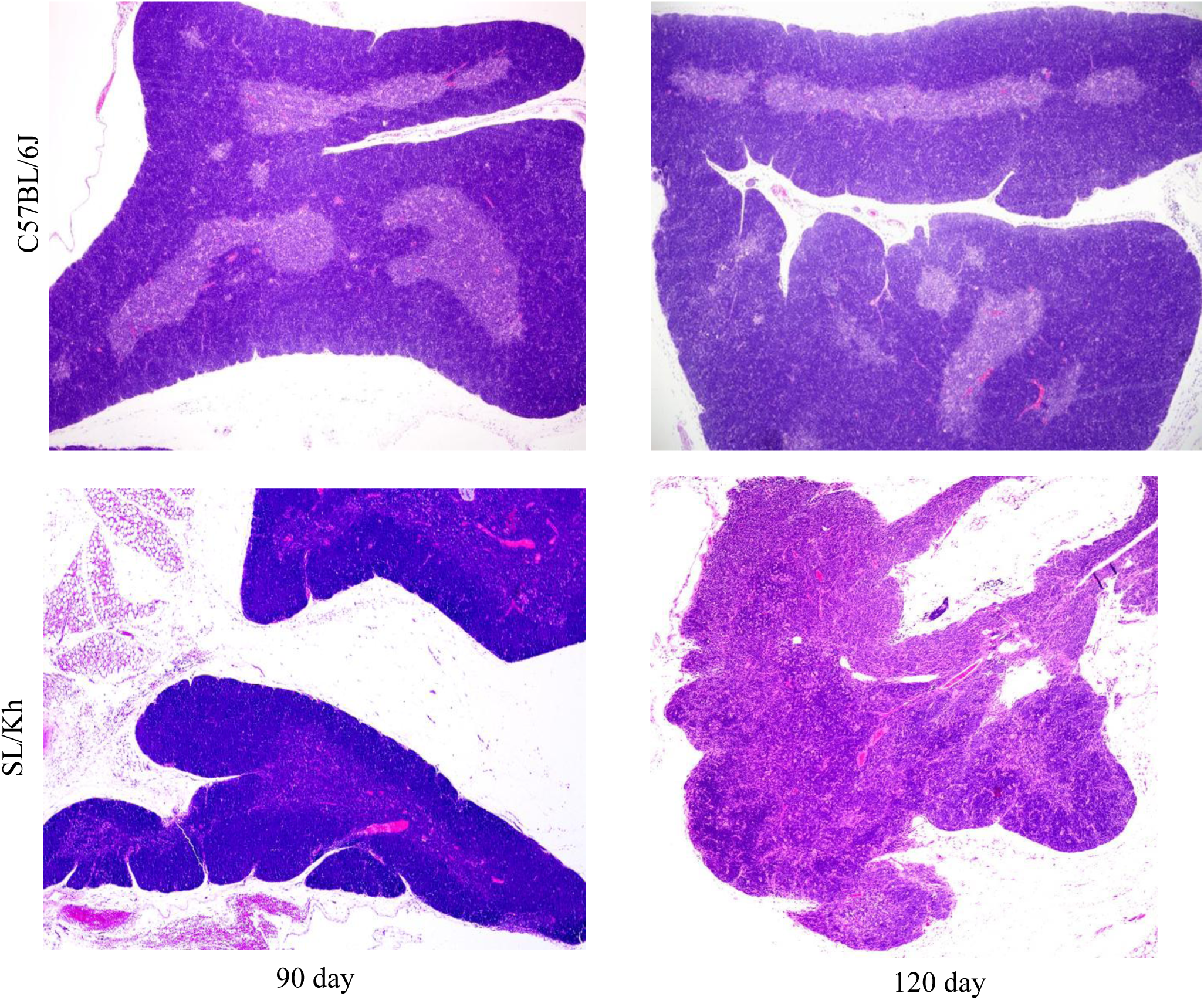
Cortical atrophy of the SL/Kh thymus. A dramatic cortical atrophy of the SL/Kh thymus can be observed when compared to C57BL/6J thymus at 90 and 120 days of age (40X). By 120 days, the normal architecture of the SL/Kh thymus is completely obliterated.

Immunohistochemical staining for two T-cell markers, CD3 and Tdt, showed a marked reduction in the number of CD3^+^, Tdt^+^ T cells in the cortex and medulla in the 120-day-old SL/Kh thymus (Figure 2A and 2B respectively). In contrast, immunohistochemical staining for Pax5, a B-cell marker, showed a dramatic increase in the number of Pax5^+^ B cells in the 120-day-old SL/Kh thymus (Figure 3). In some animals, B cell proliferation appeared to be focal or multifocal (Supplementary Figure 1), while in others a diffuse proliferation of Pax5^+^ B cells could be seen to fill nearly the entire thymus (Figure 3). At 120 days of age, there were no grossly visible thymic or mediastinal tumor masses and the spleen and lymph nodes were not enlarged. In addition, the bone marrow was histologically normal (data not shown).

**Figure 2.**
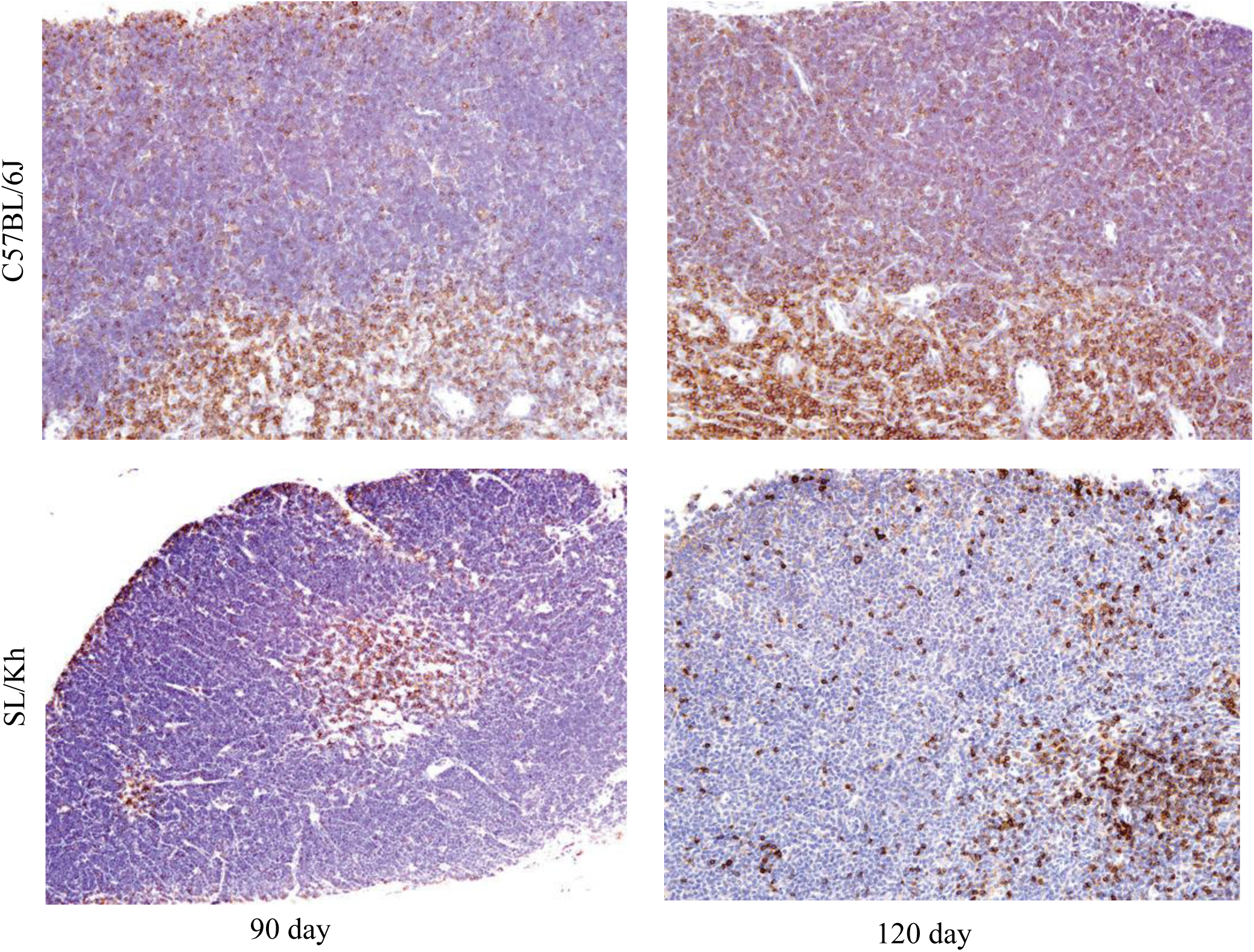
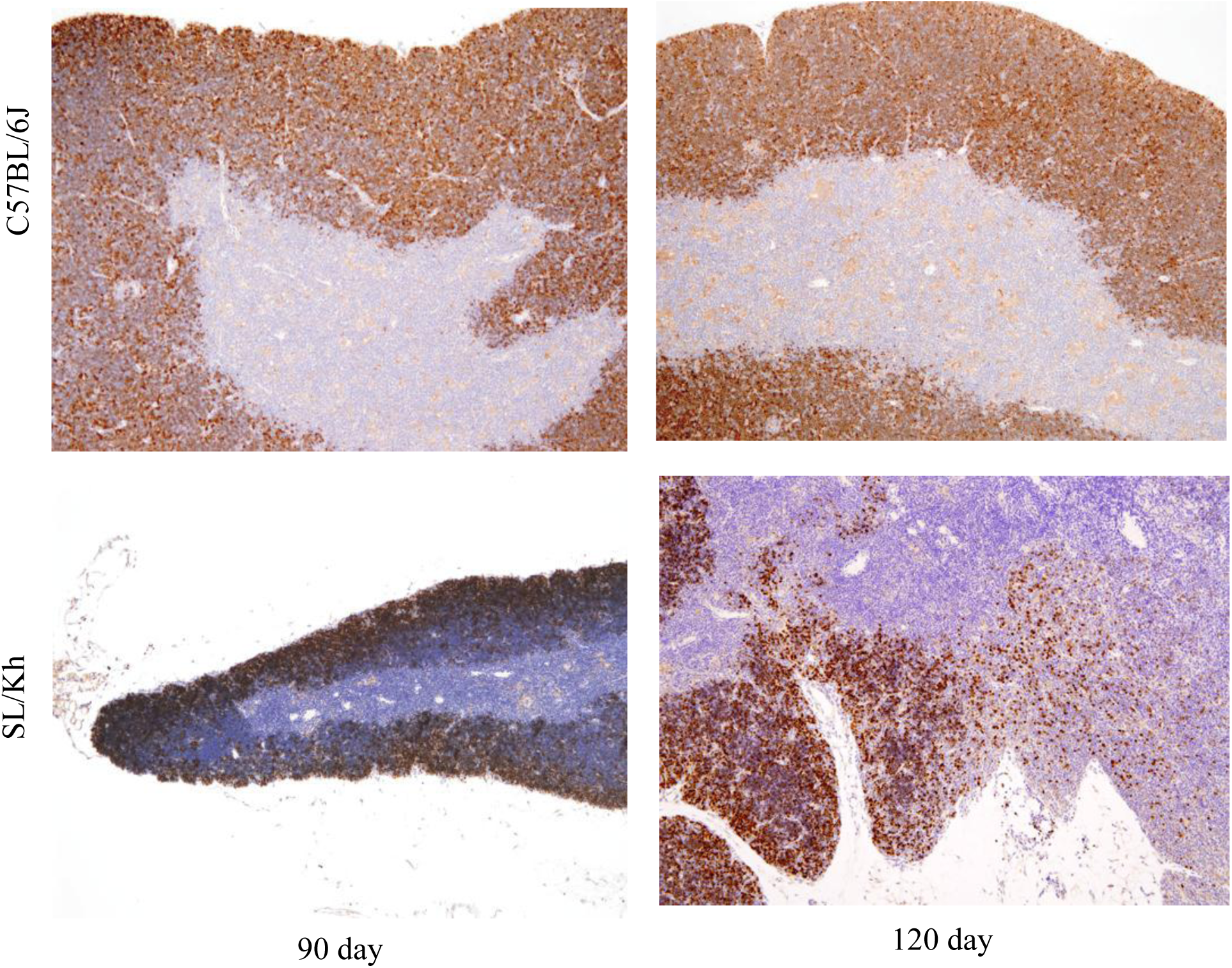
Immunohistochemical staining shows a marked reduction in the number of CD3^+^, Tdt^+^ T cells in the cortex and medulla in the 120-day-old SL/Kh thymus (A) Immunohistochemical staining for CD3, a general marker of T cells, on sections of the thymus from 90- and 120-day-old SL/Kh and control C57BL/6J mice (40X). The SL/Kh thymus is not normal and there appears to be a loss in CD3^+^ T cells in both the cortex and the medulla. (B) Immunohistochemical staining for Tdt, a marker of early T cells, on sections of the thymus from 90-(40X) and 120-day-old (100X) SL/Kh and control C57BL/6J mice (40X). While the SL/Kh thymus is already undergoing severe atrophy by 90 days, there is a clear loss of Tdt^+^ cells in the cortex of the thymus when compared to the C57BL/6J thymus.

**Figure 3.**
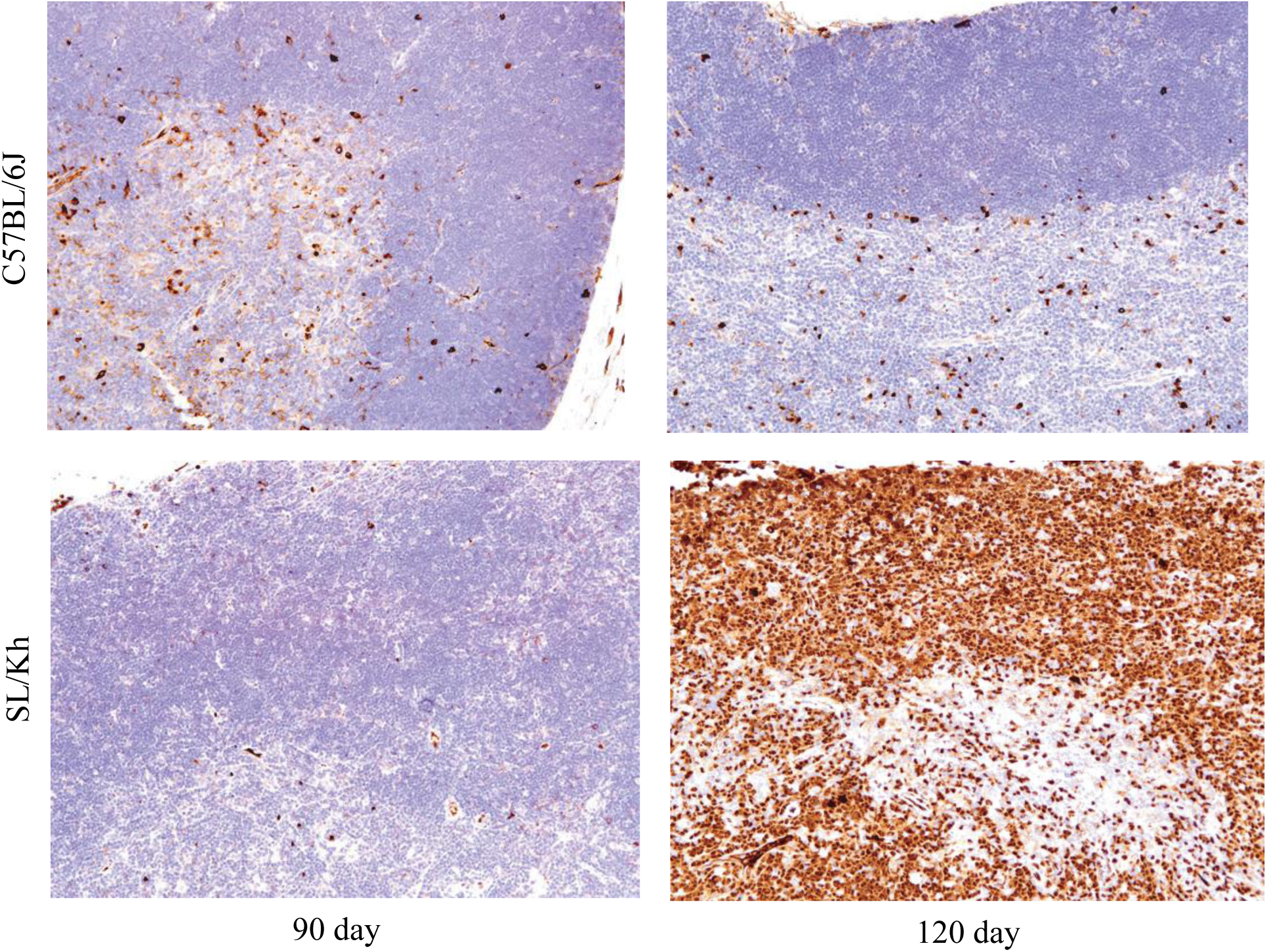
Immunohistochemical staining shows a dramatic increase in the number of Pax5^+^ B cells in the thymus of 120-day-old SL/Kh mice. Immunohistochemical staining for Pax5, a marker of early B cells, on sections of the thymus from SL/Kh and control C57BL/6J mice at 90 and 120 days of age. By120 days, the SL/Kh thymus is completely filled with Pax5^+^ B cells (200X).

Once the mice became clinically ill they were all diagnosed with pre-B lymphoma, which had obliterated the normal thymus and spread to other tissues. Among seven mice characterized in detail, three had metastases that spread to the splenic periarteriolar lymphoid sheath (PALS), lymph node paracortical high endothelial venules (HEVs) and medullary cords. The metastatic lymphoma cells were all PAX5^+^. We also observed increased numbers of PAX5^+^ B cells in areas of the spleen and lymph nodes where B cells normally reside. Furthermore, all of the 40 mice that were aged and became sick or died between 135-190 days of age showed clinical signs of lethargy, enlarged abdomens, and palpable lymph nodes. A few mice also had hind leg paralysis. In addition, the mice usually had thymic enlargement and/or mediastinal tumor masses, large spleens and enlargement of many superficial and internal lymph nodes. Histologically, the vast majority of lymphomas appeared morphologically similar.

They were composed of round to pleomorphic lymphoblasts with round nuclei, prominent nucleoli and scant cytoplasm (Figure 4A). Mitotic figures occurred in moderate numbers. The presence of prominent macrophages with cell debris (the starry sky effect) was common in thymic masses and other tissues. Metastases to the bone marrow were seen in more than half of the mice, and in the liver in most mice.

**Figure 4.**
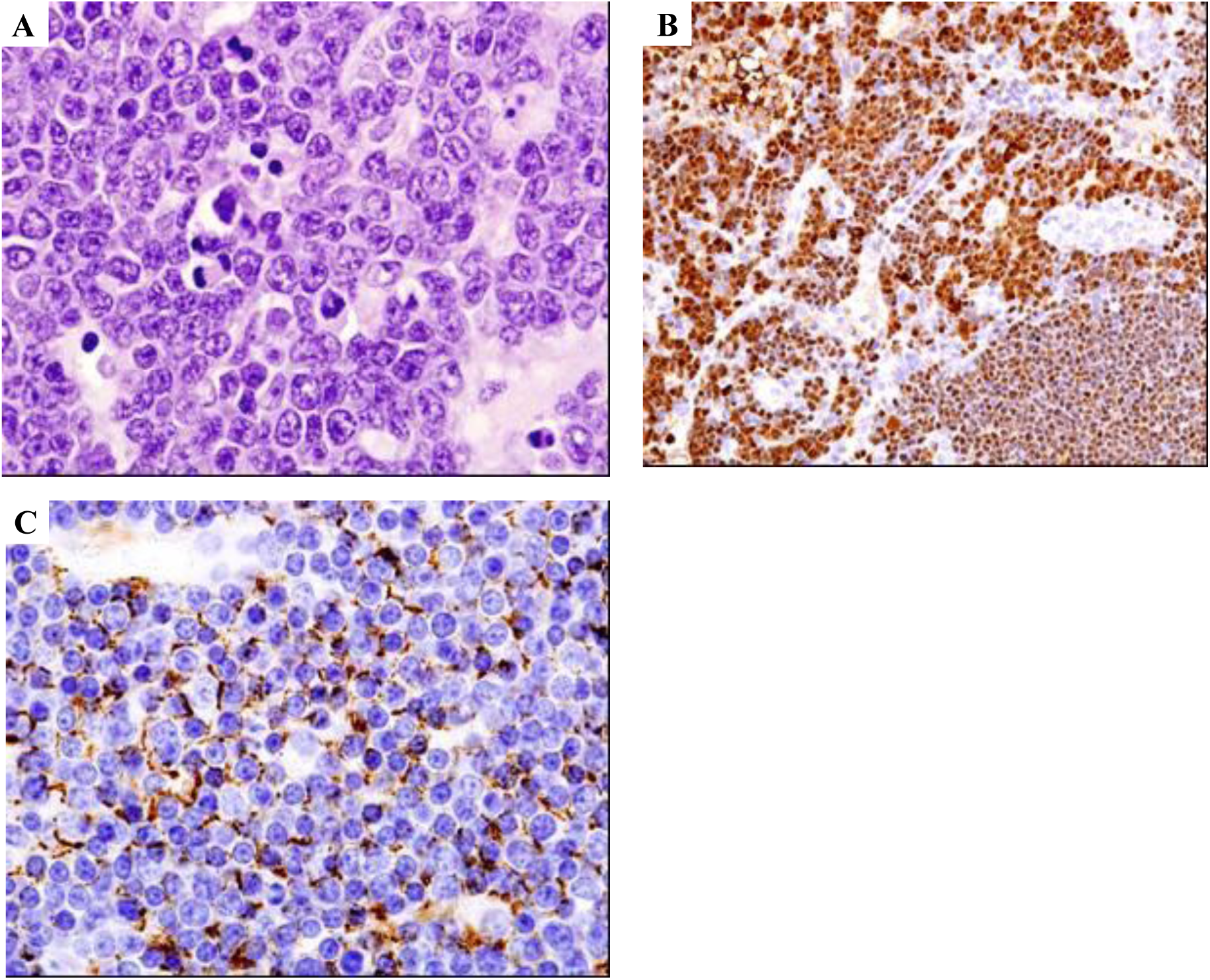
Histological analysis of SL/Kh lymphomas. (A) SL/Kh lymphomas are composed of round to pleomorphic lymphoblasts with round nuclei, prominent nucleoli and scant cytoplasm (1000X magnification). (B) Lymphomas diffusely express the nuclear B-cell marker, Pax5 (400X). Pax5^+^ lymphoma cells are shown in the top left and in the smaller normal lymphocytes in the lower right. (C) CD45R^+^ (B220), a marker of B cells, is expressed on cell membrane of lymphoma cells (X1000).

Lymphoma infiltration of the spinal meninges was also seen in some mice; some of which had clinical signs of paralysis, and others that did not. Lymphoblasts in most cases diffusely expressed PAX5 (Figure 4B) and sometimes CD45R (Figure 4C), but not CD3 or Tdt (not shown). The thymuses of all mice over 120 days were never histologically normal. Usually a lymphoma mass was seen and a few mice had early lymphomas with thymic atrophy in other lobes.

A few atypical lymphoma cases with prominent histiocytes among the tumor lymphoblasts were also observed. These cases were diagnosed as histiocyte-rich lymphomas. No cases arose from bone marrow or tissues other than the thymus, although mice were allowed to develop clinical signs after 120 days and disease dissemination; thus it was possible that some cases may have arisen in tissues other than the thymus after 120 days of age. Southern blots were performed on DNA of tumor tissues from lymphomatous mice. All tumor cells had clonal IgH rearrangements (Supplementary Figure 2), but no rearrangements in the lambda light chain or TCRβ chain, as previously reported (data not shown) ^11^. The pathological data convincingly demonstrates that the thymus is the origin of SL/Kh lymphoma.

Novel phenotypes observed in SL/Kh mice were glomerulonephritis and gal bladder hyalinosis (Ym1/Chi3l3 expression), and were seen in a majority of the mice with lymphoma (Supplementary Figure 3A). Five mice had severe diffuse mineralization of the myocardium, but mineralization in other tissues was not observed (Supplementary Figure 3B and data not shown). These potentially could be the result of host genetic factors as the glomerulonephritis and gall bladder hyalinosis was seen in a majority of animals, but it is possible that these phenotypes are the direct result of lymphoma.

### Defining the Lymphocyte Landscape of SL/Kh mice

Studies of SL/Kh lymphomas have focused almost exclusively on the spleen, lymph nodes and bone marrow, while the thymus has largely gone unstudied. This is understandable because SL/Kh lymphomas have been characterized as early pre-B cell lymphomas, and only in rare cases have B-cell lymphomas been shown to originate in the thymus ^10, 11^. For our study it was imperative, therefore, that we examined pre-lymphomatous tissues, including the thymus, to characterize the shifting lymphocyte landscape as it relates to disease progression, guided by our histopathological analysis.

To accomplish this, single cell suspensions from the bone marrow, spleen and thymus were labeled with three color antibody cocktails designed to identify the major subsets of B and T cells by direct immunofluorescent staining using flow cytometric analysis. The percentage of each cell population was then determined for SL/Kh bone marrow, spleen and thymus, and the significance of these percentages relative to the B6 control was determined using a student’s t-test (Figure 5A-F, Supplementary Figure 4A-4O).

**Figure 5.**
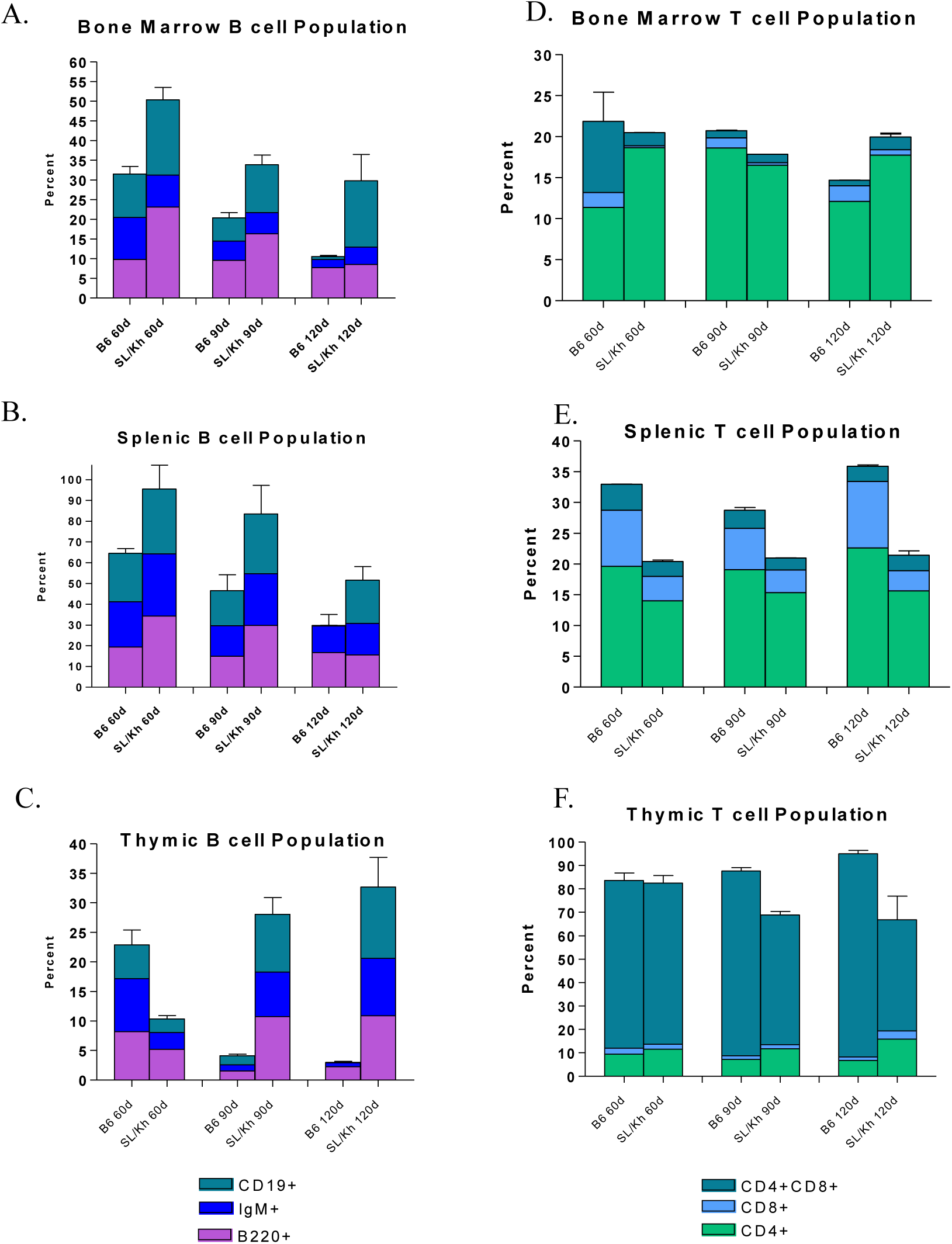
The lymphocyte landscape of SL/Kh versus control C57BL/6J mice at 60, 90 and 120 days of age. Graphs were compiled from FACS data and plotted according to the percentage of cells that stained positive for the indicated cell surface markers. Statistical analysis was performed using a student *t-test*^58^. B cell populations that were significantly different (p<0.01) between SL/Kh and control C57BL/6J mice for bone marrow were as follows: (A) CD19 at 90 and 120 days, IgM at 60 and 120 days, and B220 at 60 days for spleen; (B) CD19 at 120 days, and B220 at 60 and 90 days for thymus; (C) CD19 at 90 and 120 days, IgM at 90 and 120 days, and B220 at 90 and 120 days for spleen. T cell populations that were significantly (p<0.01) different between SL/Kh and control C57BL/6J mice for bone marrow were as follows: (D) CD8 at 60, 90 and 120 days and CD4 at 60 and 120 days for spleen; (E) CD4, CD8 at 60 days, CD8 at 60, 90 and 120 days, and CD4 at 60 days for thymus; (F) CD4, CD8 at 90 and 120 days, CD8 at 120 days, and CD4 at 120 days.

SL/Kh mice had significantly more B cells in the bone marrow (Figure 5A) and spleen (Figure 5B) at 60, 90 and 120 days of age compared to B6 mice; however the total population of B cells decreased with age for both strains of mice. This increased number of B cells is likely due to the transient pre-B expansion that has been previously reported for SL/Kh mice ^12^. As one would expect, the B cell population drastically decreases with age in the thymus of B6 mice (Figure 5C). In contrast, the SL/Kh thymus shows a significant increase in the number of B cells with age (Figure 5C). In summary, the thymus, bone marrow and spleen of SL/Kh mice have abnormally high levels of B cells that persist until the time of onset of lymphoma development. Most unusual is the rapid accumulation of B cells in the thymus, a tissue that is primarily composed of T cells. Of note, these B cells are mostly CD25^-^ (described below), which would indicate that the lymphoma is blocked at the pro/pre-B stage of development, not the pre-B stage of development as previously reported.

The total T cell population in the bone marrow of B6 mice also decreases with age (Figure 5D), whereas in the spleen and thymus, T cell levels persist at relatively high levels (Figure 5E and 5F). The total T cell populations in the SL/Kh bone marrow and spleen remains relatively constant with age; however the total levels of T cells in these tissues is significantly less than in B6 mice (Figure 5D and 5E). This suggests that these tissues have an abnormally low baseline level of T cells, but retain the ability to regulate homeostasis. The T cell populations in the thymus of B6 and SL/Kh mice were similar at 60 days of age (Figure 5F). By 90 and 120 days, however, the B6 thymus retains a relatively high and constant level of T cells, whereas the SL/Kh thymus shows an overall decrease in T cells specific to the CD4/CD8 double positive population (Figure 5F). Of note, there is a significant increase in the CD4 single positive cells versus the same population in B6 thymus (Figure 1F). In summary, the SL/Kh thymus shows a rapid increase in B cells that correlates with a simultaneous loss of T cells.

### Block in early T lymphocyte development in the SL/Kh thymus

To determine whether there was a defect in T cell development that could explain the rapid increase in B cell number in the thymus of SL/Kh mice, single cell suspensions were made from the thymus of 45-day-old SL/Kh and control B6 mice, and labeled with CD4, CD8, CD44 and CD25 antibodies ^21^. The CD4, CD8 double negative (DN) population was then sorted and analyzed for expression of CD44 and CD25. This sorting scheme made it possible to determine the nature of the mature and immature T cell populations in the thymus of SL/Kh mice based on known genetic checkpoints (Figure 6) ^21, 22^. At 45 days of age, the SL/Kh thymus had significantly fewer CD4, CD8 double positive cells, and a significantly higher DN population, than B6 control mice (Figure 7A and 7B). In addition, sorting of the DN cells also revealed a significant decrease in the double positive CD44, CD25 population and single positive CD25 population, but a significant increase in the single positive CD44 population (Figure 7C and 7D). SL/Kh thymic T cells thus appear to be blocked at the CD4^-^, CD8^-^, CD44^+^, CD25^-^ stage of development (Figures 5, 6, and 7). Interestingly, it is this population that has the capability of differentiating into pre/pro-B, dendritic and natural killer (NK) cells (Figure 6). No difference in TUNEL staining was observed between SL/Kh and control B6 thymuses, suggesting that the reduced number of T cells in the SL/Kh thymus is not due to apoptosis but rather from a block in development (data not shown). One possible cause of the developmental defects observed in the SL/Kh thymus is a defect in Notch1 expression. Notch1 is a key regulator of T cell development ^23–25^.

**Figure 6.**
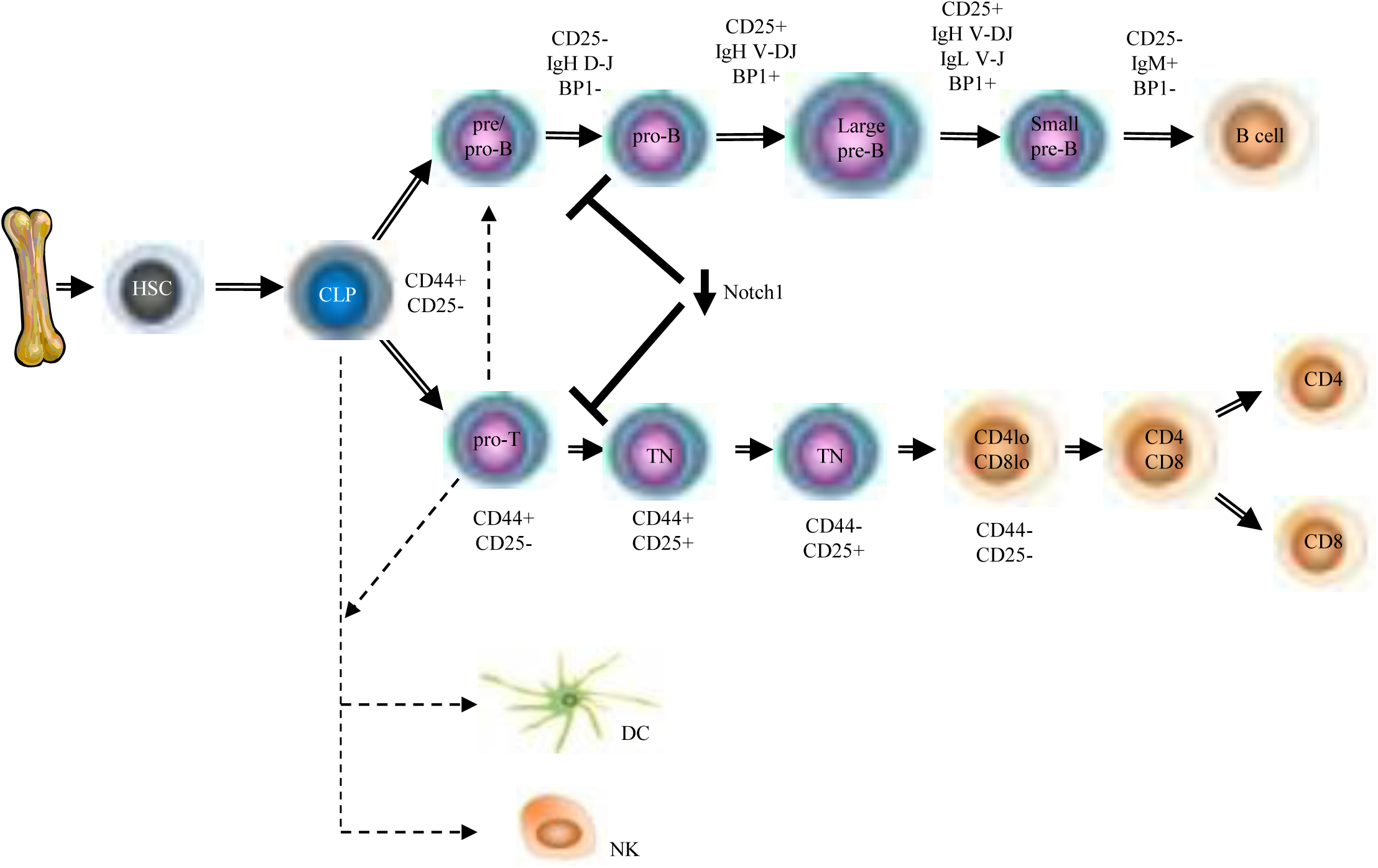
Schematic diagram of mouse B- and T-cell development. Hematopoietic stem cells (HSCs) are produced in the bone marrow. The common lymphoid progenitor (CLP) is derived from HSCs and can differentiate into B cells, T cells, dendritic cells (DC) and natural killer (NK) cells. The exact stage of hematopoietic cell development can be delineated through the analysis of the expression of various cell surface markers. Cells that have the surface phenotype of CD4^-^, CD8^-^, CD44^+^, CD25^-^ are thought to be CLPs. When Notch1 expression is down-regulated in the thymus there is a differentiation block at the pre/pro-B and pro-T cell stage of development.

**Figure 7.**
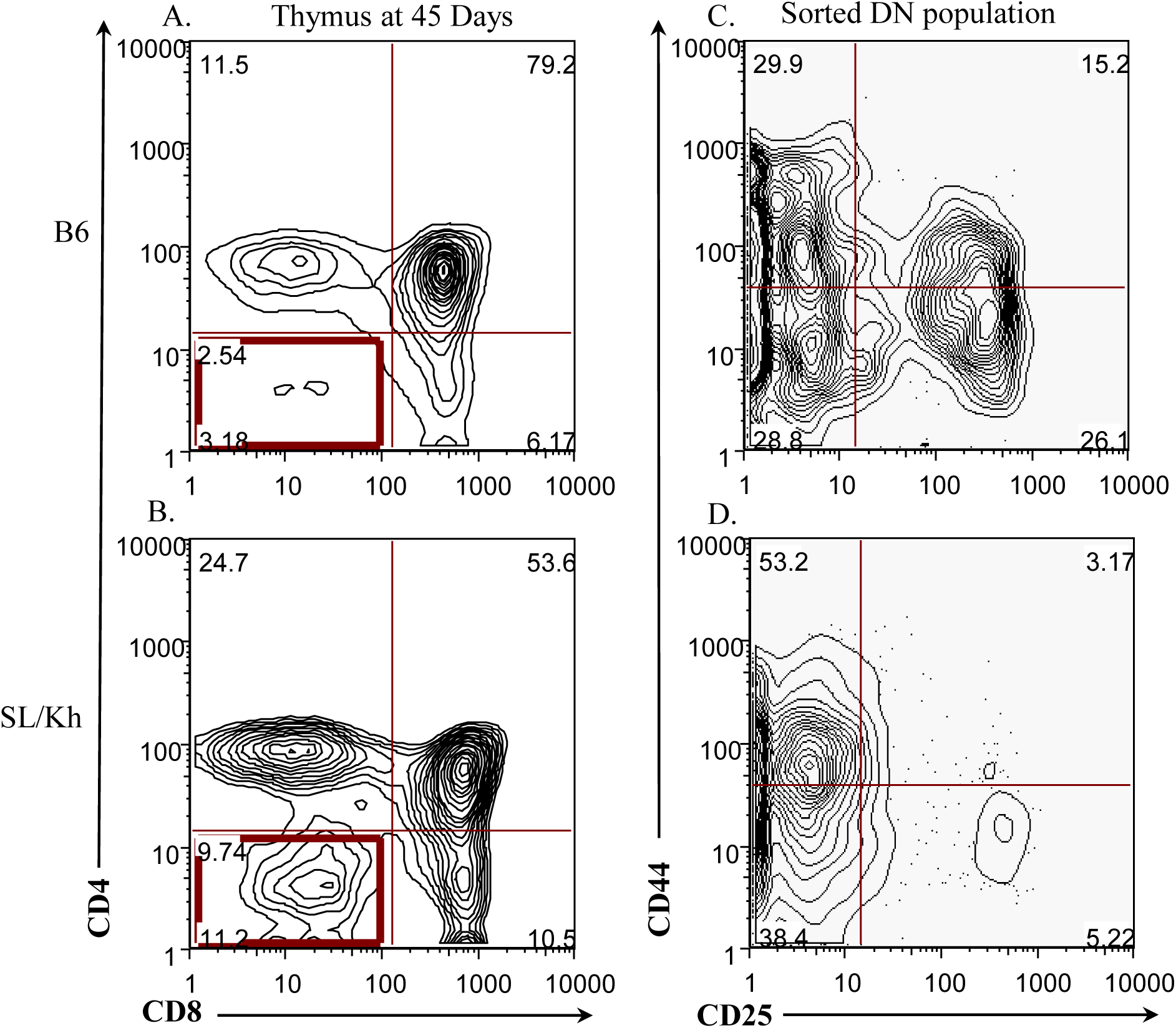
A block in early T cell development in the SL/Kh thymus. (A, B) At 45 days of age the SL/Kh thymus has significantly fewer CD4, CD8 double positive T cells, and a significantly higher number of double negative cells, than the C57BL/6J control thymus. (C, D) Sorting of the double negative population revealed a significant decrease in the double positive CD44, CD25 population and single positive CD25 population, but a significant increase in the single positive CD44 population, in the SL/Kh thymus relative the C57BL/6J control thymus. A red box indications the double negative population.

When its expression is inactivated in bone marrow precursors, thymic T cells become blocked at an early stage of T cell development with a concomitant increase in the proliferation of an early B cell population (Figure 6) ^26^, just like what we observed in thymus of SL/Kh mice. To determine whether a defect in Notch1 expression could account for the T cell defects observed in SL/Kh mice, we measured the levels of Notch1 expression in whole thymus and sorted T cell populations isolated from 45-day-old SL/Kh and control B6 mice using RT-PCR analysis. Interestingly, both the whole thymus and sorted CD4^-^, CD8^-^, CD44^+^, CD25^-^ cell population from SL/Kh mice showed a significant down-regulation of Notch1 expression (Figure 8). A trivial explanation for this result could be the increased ratio of B to T cells observed in the SL/Kh thymus. This is unlikely to be the explanation, however, since the thymus and sorted CD4^-^, CD8^-^, CD44^+^, CD25^-^ cells were obtained from 45-day-old SL/Kh mice, well before these mice exhibit a thymic phenotype. In addition, the sorted CD4^-^, CD8^-^, CD44^+^, CD25^-^ subpopulation showed the same down-regulation of Notch1 expression seen in whole thymus, indicating that this defect in Notch1 expression is intrinsic to the T cells and not solely because the SL/Kh thymus is undergoing atrophy or shifting the lymphocyte lineage from T- to B-cells.

**Figure 8.**
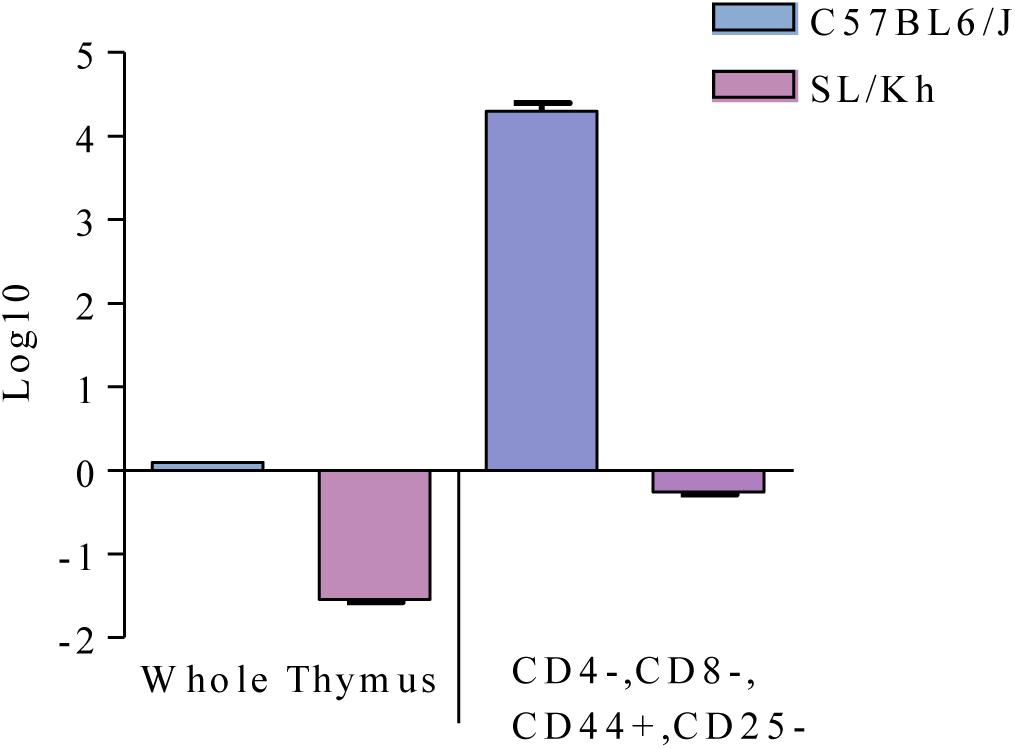
Defective Notch1 expression in the thymus of SL/Kh mice. Levels of Notch1 expression in whole thymus and sorted T cell populations from 45-day-old SL/Kh and control C57BL/6J thymus using RT-PCR analysis. Data is normalized to Notch1 expression in the C57BL/6J whole thymus. The statistical difference in Notch1 expression between SL/Kh and C57BL/6J control thymus is p<0.003 for both groups.

### Identification of common retroviral insertion sites in SL/Kh lymphomas

Previous insertional mutagenesis profiling of SL/Kh lymphomas identified *Stat5a*, *Stat5b*, *Zfp521* (also called *Evi3*), *Myc* and *Nmyc* as genes that are insertionally mutated in SL/Kh lymphomas^13^. Here, we sought to extend this list and identify additional genes involved in SL/Kh lymphomagenesis. In our experiments, we identified 42 genes that were insertionally mutated in at least 2 of 40 SL/Kh lymphomas analyzed (Supplementary Table 1). Genes that are insertionally mutated in multiple lymphomas are believed to have the highest relevance to disease onset and progression. As previously reported, we identified insertions in *Stat5a*, *Stat5b*, *Myc* and *Nmyc*, but only in a single lymphoma each. We also identified integrations in *Zfp521* in lymphomas from 5 different animals. Interestingly, the 42 insertionally mutated genes clustered in two main pathways with about 20% of the genes involved in the RAS/MAPK/ERK pathway, and another 20% involved in the JAK/STAT and B cell development pathways (Figure 9).

**Figure 9.**
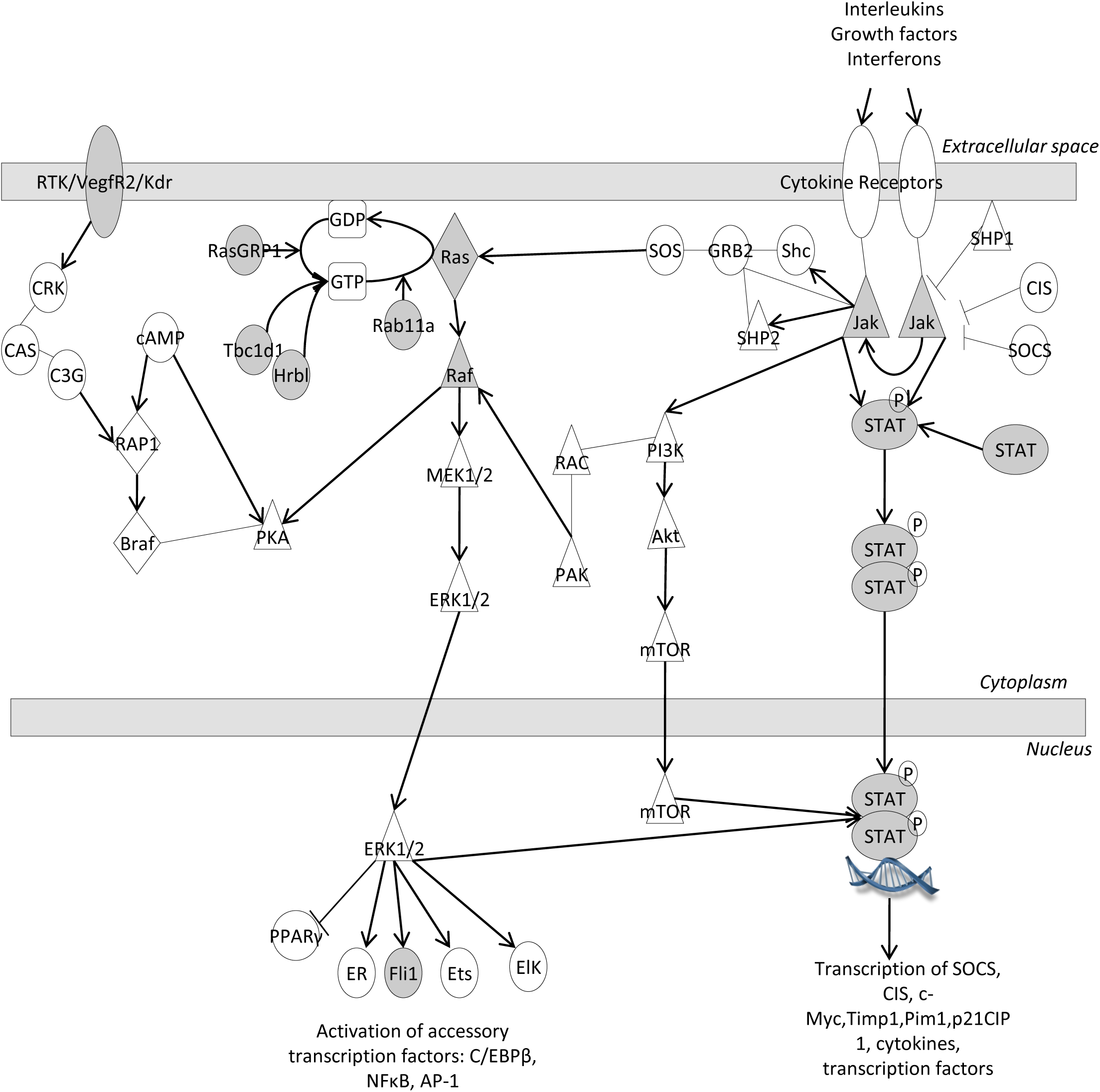
Pathway diagram illustrating some of the genes known to function in the RAS/MAPK/ERK and JAK/STAT signaling pathways. Gray-shaded characters are genes that were found to be insertionally mutated in two or more SL/Kh lymphomas. While STAT was only insertionally mutated in one lymphoma it is also highlighted in gray because it was shown to be insertionally mutated in multiple SL/Kh lymphomas in previous studies ^59, 60^.

Both of these pathways are known to signal in normal B-cell development ^27, 28^ and are misregulated in B-cell lymphomas ^29, 30^. The fact that these genes were enriched among those insertionally mutated in SL/Kh lymphomas is not surprising because the cell population that is rapidly expanding in preleukemic SL/Kh mice is early B cells, which thus represent the expanding pool of cells for insertional mutagenesis.

## Discussion

We demonstrate here for the first time that SL/Kh pre-B lymphomas originate in the thymus and not the lymph nodes, spleen or bone marrow as previous studies have implied. We also describe a novel thymic defect in SL/Kh mice that results in thymic atrophy, a block in early T cell development with a concomitant reduction in the number of thymic T cells as the animal ages, and an expansion in the thymic pro/pre-B cell population. It is this genetic defect(s), which increases the likelihood that retroviral insertional mutagenesis will occur in thymic B cells, that we believe makes SL/Kh mice so uniquely susceptible to thymic pre-B lymphomas.

Although we did not uncover the exact cause of this genetic defect(s) we show that it is associated with decreased Notch1 expression. Studies using CD3ε transgenic mice ^31^, Notch1-deficient mice ^32^ or TCRβ-deficient mice ^33^ all show that the number of thymic B cells increases significantly in association with decreasing thymic T cells. They also suggest that in the normal thymus, T cells take up most of the microenvironmental niches, which prevents efficient thymic B cell maturation and colonization. Notch1 signaling in the thymus is proposed to work through a lateral inhibition mechanism where two cells that are initially identical have different Notch1-sending (down-regulation) and Notch1-receiving (up-regulation) roles ^34, 35^. When a T cell expresses high levels of Notch1, the surrounding cells recognize the Notch1 signal and T-fated cells are directed to differentiate or proliferate as T cells. Cells with other developmental fates, such as pro/pre B cells, remain in a static state. When Notch1 signaling is downregulated, pro/pre-B cells no longer receive inhibitory Notch1 signals allowing their uncontrolled proliferation. The reduced number of T cells present in the SL/Kh thymus, combined with the 10-fold reduction in Notch1 expression in the remaining T cells, might therefore not produce enough of an inhibitory Notch1 signal to keep pro/pre-B cells in the thymus in check or stop the common lymphoid progenitor (CLP) from adopting a B cell fate, thus explaining the pre-B expansion observed in the SL/Kh thymus. In support of this hypothesis, our data show that thymic SL/Kh T cells are blocked at the CD4^-^, CD8^-^, CD44^+^, CD25^-^ stage of development, which is exactly same stage where Notch1-deficient T cell are blocked in development ^32^.

A precedent for these findings can be found in studies of another leukemia prone strain, BXH2. Like SL/Kh mice, these mice were unusual among the early leukemia prone strains in that they develop a high frequency of retrovirally induced myeloid leukemia that is uniformly fatal by one year of age. In an elegant series of genetic experiments, Turcotte et al. showed that BXH2 mice carry a missense mutation in a transcription factor of the interferon (IFN) regulatory factor (IRF) family, interferon regulatory factor 8 (*Irf8*) ^36^. The IRF family of proteins bind to the IFN-stimulated response element and regulate expression of type I IFN response genes. Mice with a targeted mutation in *Irf8* are immunodeficient and develop a syndrome similar to human chronic myelogenous leukemia^37^. The chronic period of the disease progresses to a fatal blast crisis characterized by a clonal expansion of undifferentiated cells, suggesting a novel role for *Irf8* in regulating the proliferation and differentiation of hematopoietic progenitor cells. These results suggest a two-step model for leukemia induction in BXH2 mice, in which inactivation of *Irf8* predisposes to myeloproliferation and immunodeficiency. This in turn leads to increased levels of retroviral replication and subsequent insertional mutagenesis of the expanded myeloid cell pool, which eventually leads to myeloid leukemogenesis.

It is well known that the thymus is the major site of T cell differentiation but for the last ten years it has been increasingly appreciated that the thymus also houses a resident population of B cells. Since all hematopoietic progenitors arise in the bone marrow, a candidate common lymphoid progenitor (CLP) population characterized by surface expression of CD44^+^,CD25^-^ has been proposed to be the source of the resident B cells in the thymus ^38^. It has also been shown that cells which are CD4^-^, CD8^-^, CD44^+^,CD25^-^ have the potential to differentiate into T, B, NK cells and dendritic cells ^38–40^. The thymus contains all of the molecular signals necessary to support the growth of early B cells and it is thought that thymic B lymphopoiesis occurs concomitantly with migration from the cortex to the medulla of the thymus, identical to T lymphopoiesis ^41^. Since all thymic progenitor cells are derived from the bone marrow (BM) it is likely that the seeding cell/CLP migrates from the BM to the thymus. It is thus reasonable to conclude that it is the resident B cell population derived from CLP migration from the BM to the thymus that is the source of cells for lymphomagenesis in the SL/Kh mice.

When we quantitated the B cell populations present in the BM, spleen and thymus of SL/Kh mice throughout development, we found that the B cell population decreases in the BM and spleen as the animal ages, but increases dramatically in the thymus. There are two possibilities that could account for this expansion. One possibility is that this results from an expansion of the resident thymic pro/pre-B cell population. Another is that the CLP is forced to adopt a B cell fate instead of the normal T cell fate. Data from thymectomy studies suggest that the BM CLP pool is responsible for the expanded B cell population observed in SL/Kh mice. Lu and colleagues have reported that thymectomized SL/Kh (Tx-SL/Kh) mice develop lymphomas with a pro/pre B phenotype and a latency that is doubled compared to intact SL/Kh mice ^42^. Thymectomy removes the resident B cells in the thymus; however, the animals still develop pro/pre-B lymphomas, suggesting that the CLP derived from the BM is the likely source of lymphomagenic potential. It is interesting that Tx-SL/Kh mice presented with the same disease but a doubled latency. In future studies it will be important to determine whether the cervical thymi or some other organ site provides the necessary niche for lymphomagenesis to occur in Tx-SL/Kh mice.

Primary mediastinal (thymic) large B cell lymphoma (MLBCL) is a human lymphoma that is derived from thymic B cells. MLBCL was first described in the 1980’s but it took 14 years to be accepted by pathologists and clinicians as a distinct subtype of large diffuse B-cell lymphoma because of its unique histomorphological, immunological and genetic profile ^43–45^. The discovery of normal B cells in the thymus that share immunophenotypical similarities with MLBCL has led to the accepted hypothesis that thymic B cells are the precursors of MLBCL ^46–48^. Like SL/Kh thymic pre-B-lymphoma, human MLBCL is a highly aggressive tumor that arises in the thymus and affects females at a slightly higher incidence than males. MLBCL cells also express B-cell lineage specific surface molecules and lack surface immunoglobulin (Ig) expression ^10^. With a few exceptions, MLBCL also shows constitutive expression of the immunoglobulin-associated molecule CD79a ^49^, which is thought to hold the B cells in an immature state^50^. Patients with MLBCL rarely present with invasion to the bone marrow, similar to what has been observed for SL/Kh lymphomas where bone marrow involvement is not a defining aspect of the disease in the majority of cases. There are thus many similarities between SL/Kh lymphoma and human MLBCL.

Similar to BXH2 myeloid leukemias, we envision that SL/Kh pre-B lymphomas occur through a two-step process that involves expansion of the pre-B cell pool in the thymus combined with subsequent retroviral insertional mutagenesis of the expanded B cell pool. In previous studies, *Stat5a*, *Stat5b*, *Evi3*, *Myc* and *Nmyc* were identified as genes that are activated by proviral integration in SL/Kh lymphomas ^13^. In our studies we extended this list and identified additional genes that are predicted to be involved in SL/Kh lymphomagenesis. Among these genes, 20% are predicted to function in the RAS/MAPK/ERK pathway, while another 20% are predicted to function in JAK/STAT signaling and B cell development pathways. Both of these pathways signal in normal B cell development ^27, 28^, and both are misregulated in human B lymphomas ^29, 30^. That we hit genes involved in pathways of early B cell development and lymphomagenesis is not surprising because this is the cell population that is rapidly expanding in the SL/Kh thymus as the animal ages and is thus the natural target for retroviral insertional mutagenesis.

The interleukin-7 (IL-7) receptor is the only interleukin receptor known to be essential for early B cell development. Mice deficient for *Il7rα* ^51^, the shared γ_c_ receptor (*IL2rg*)^52^ or *Jak3* ^53^ all manifest a block at the pro/pre B cell stage of development. It has been proposed that IL-7 activates Ras in B cell progenitors or that activation of Ras by some other factor permits IL7 receptor induced STAT signaling ^28^. Maximal STAT activation in turn requires MAP kinase mediated serine phosphorylation. In the case of SL/Kh lymphomas, *Ras*, *Jak*, *Stat5a*, and *Stat5b* are all activated via proviral integration, which either singly or in combination promotes downstream signaling to MAPK, resulting in uncontrolled cell proliferation. We also observed IL7, IL4 and Nras mRNA up-regulation in all SL/Kh lymphomas (data not shown). Interestingly, human MLBCL shows a transcriptional signature of constitutively activated NFκB, the JAK-STAT pathway, and several members of the interleukin-4 (IL-4)/IL-13 pathway ^54, 55^.

Inactivating mutations in suppressors of cytokine signaling (SOCS) family members have also been described for MLBCL ^56^. SOCS family members are part of a classic negative feedback loop involving JAK/STAT and Ras pathways where inhibition of SOCS leads to sustained activation of Ras ^57^. The overlap in gene expression between human MLBCL and SL/Kh pre-B lymphomas, combined with the pathological similarities, are additional reasons that SL/Kh should be considered as a model for MLBCL.

## Acknowledgements

We would like to thank Keith Rogers, Susan Rogers and the histopathology core at IMCB for helpful discussions, mouse necropsies, and tissue and slide preparations; Alice Tay and the DNA sequencing core at IMCB for help with tumor DNA sequencing; and our animal room technicians, Nicole Lim, Pearlyn Cheok and Dorothy Chen for strain maintenance and tumor monitoring. Finally, we would thank Motomi Osato for advice on FACs analysis. Support was provided by the Biomedical Research Council (BMRC), Agency for Science, Technology and Research (A*STAR), Singapore.

## Supplementary figure legends

**Supplementary Figure 1.**
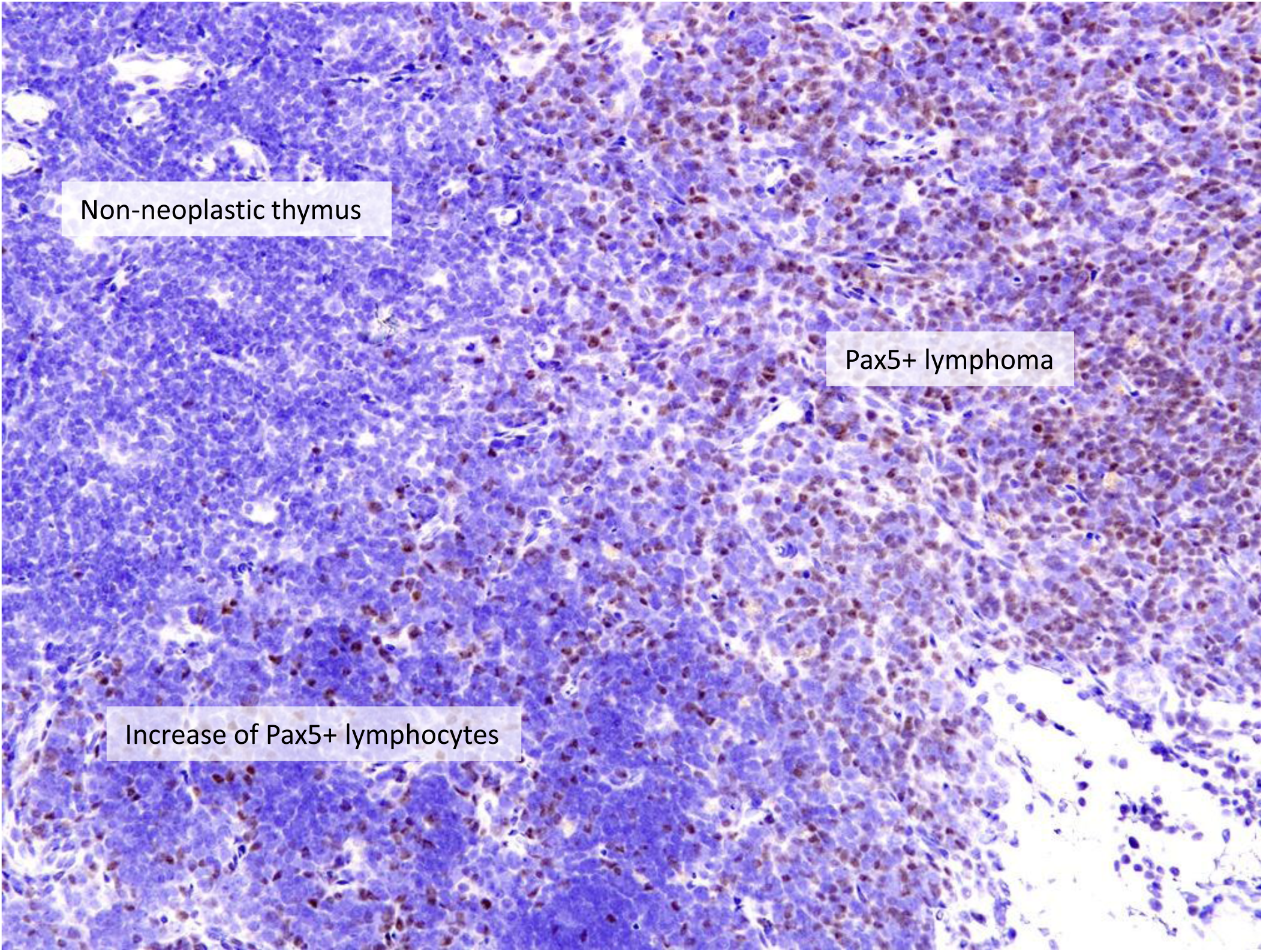
Pax5 staining of early lymphoma in the thymus of SL/Kh mice. Pax5 positive cells are indicated by brown staining (200X).

**Supplementary Figure 2.**
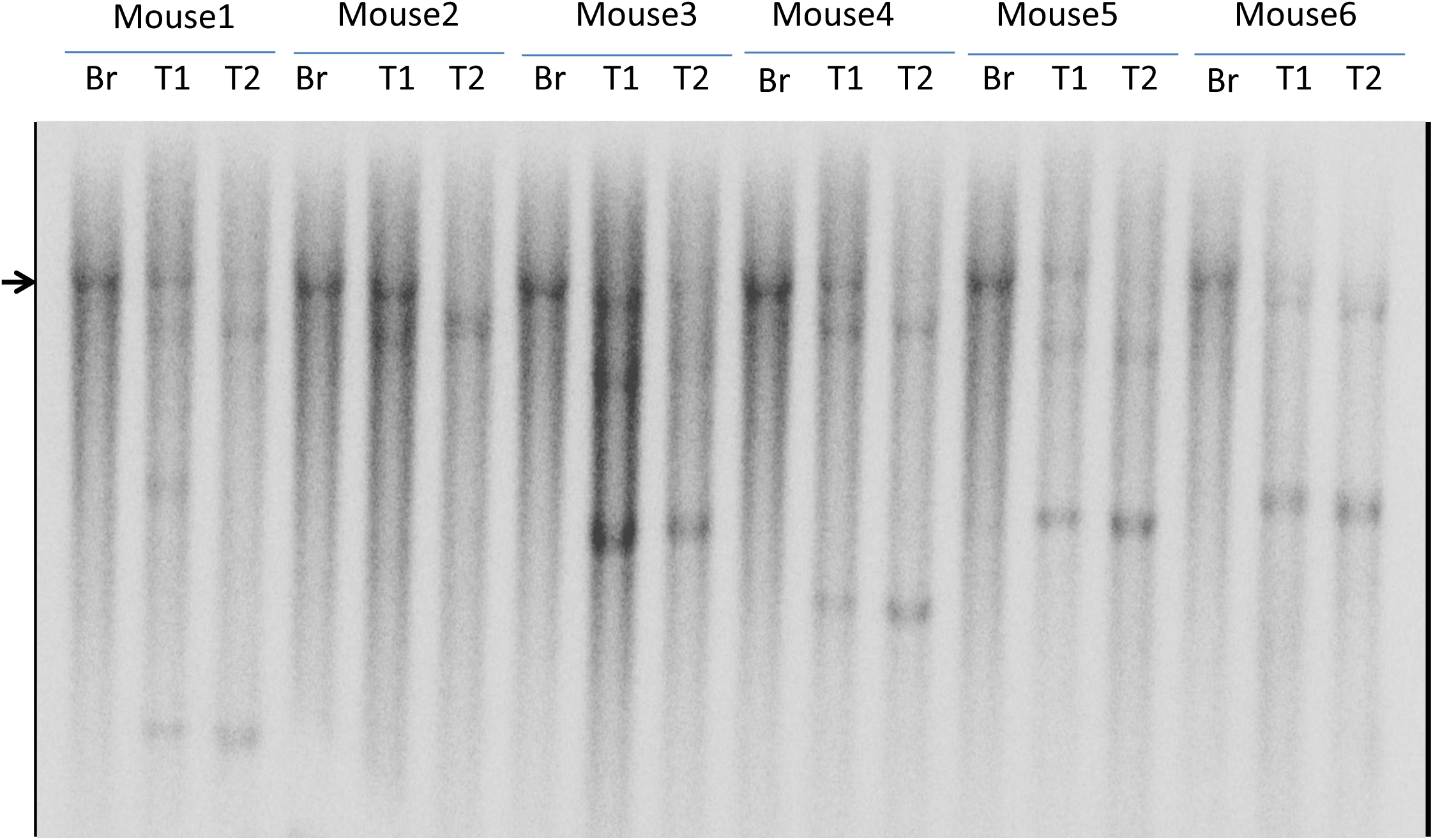
SL/Kh lymphomas have clonal IgH rearrangements. Southern blot analysis of SL/Kh lymphomas from two lymphomatous tissues from each animal (T1, T2) and control brain (Br) demonstrates that tumors have clonal IgH rearrangements. The arrow designates the un-rearranged band.

**Supplementary Figure 3.**
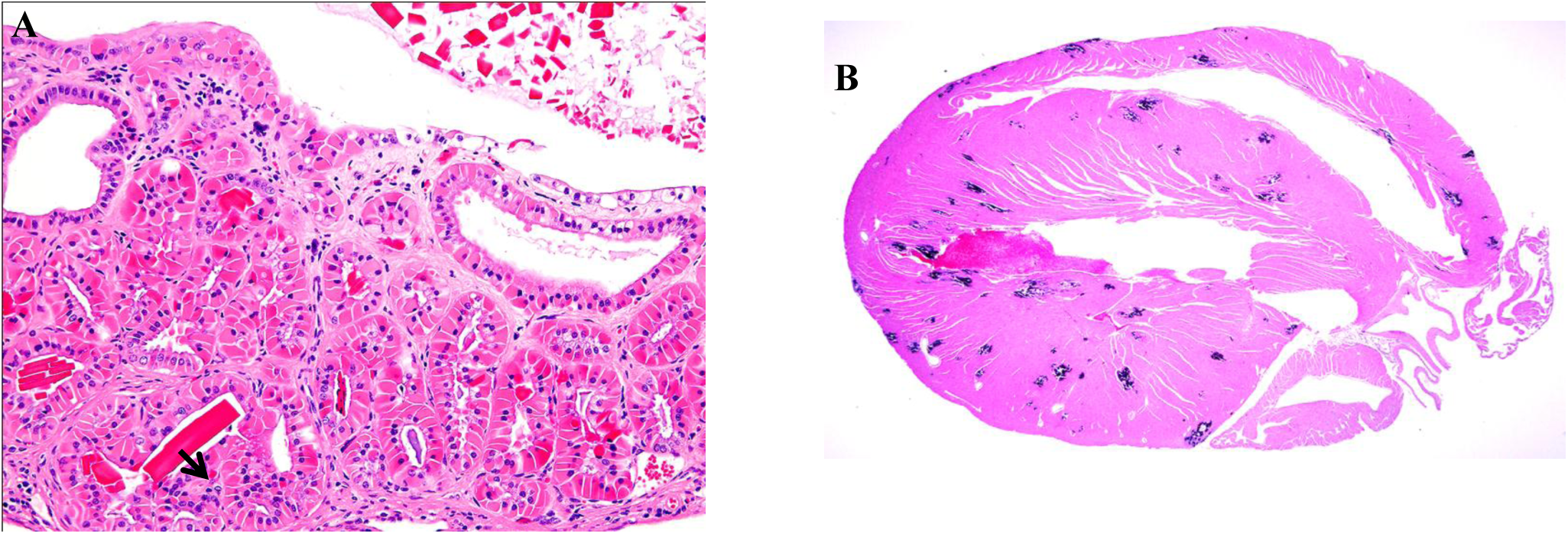
Novel pathological findings in the gall bladder and heart of SL/Kh mice. (A) H&E stained section showing hyalinosis with crystal formation in the gall bladder, X200. The arrow indicates one of the crystal formations. (B) H&E stained section showing disseminated calcification of the heart myocardium (blue foci) (20X).

**Supplementary Figure 4.**
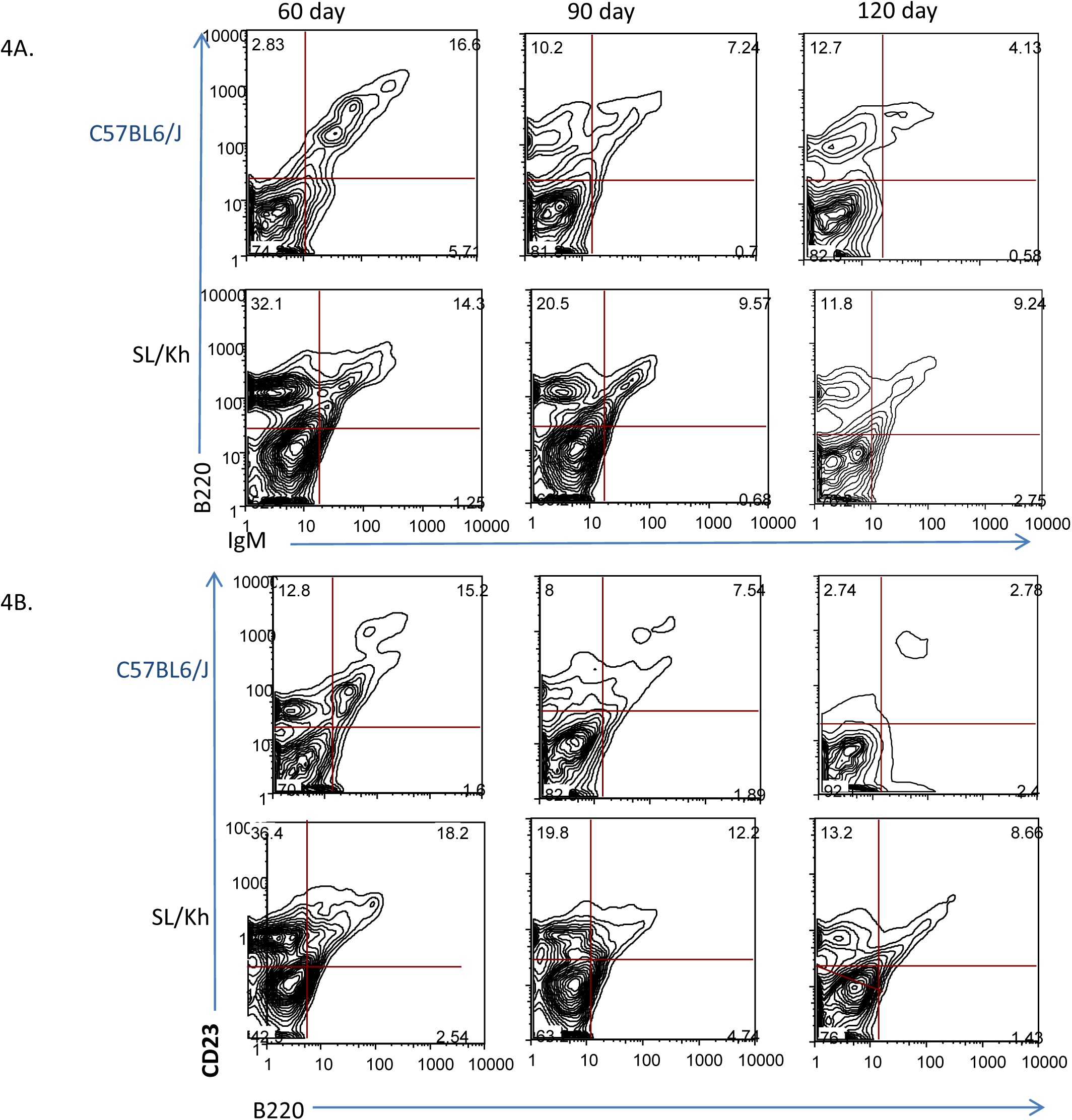
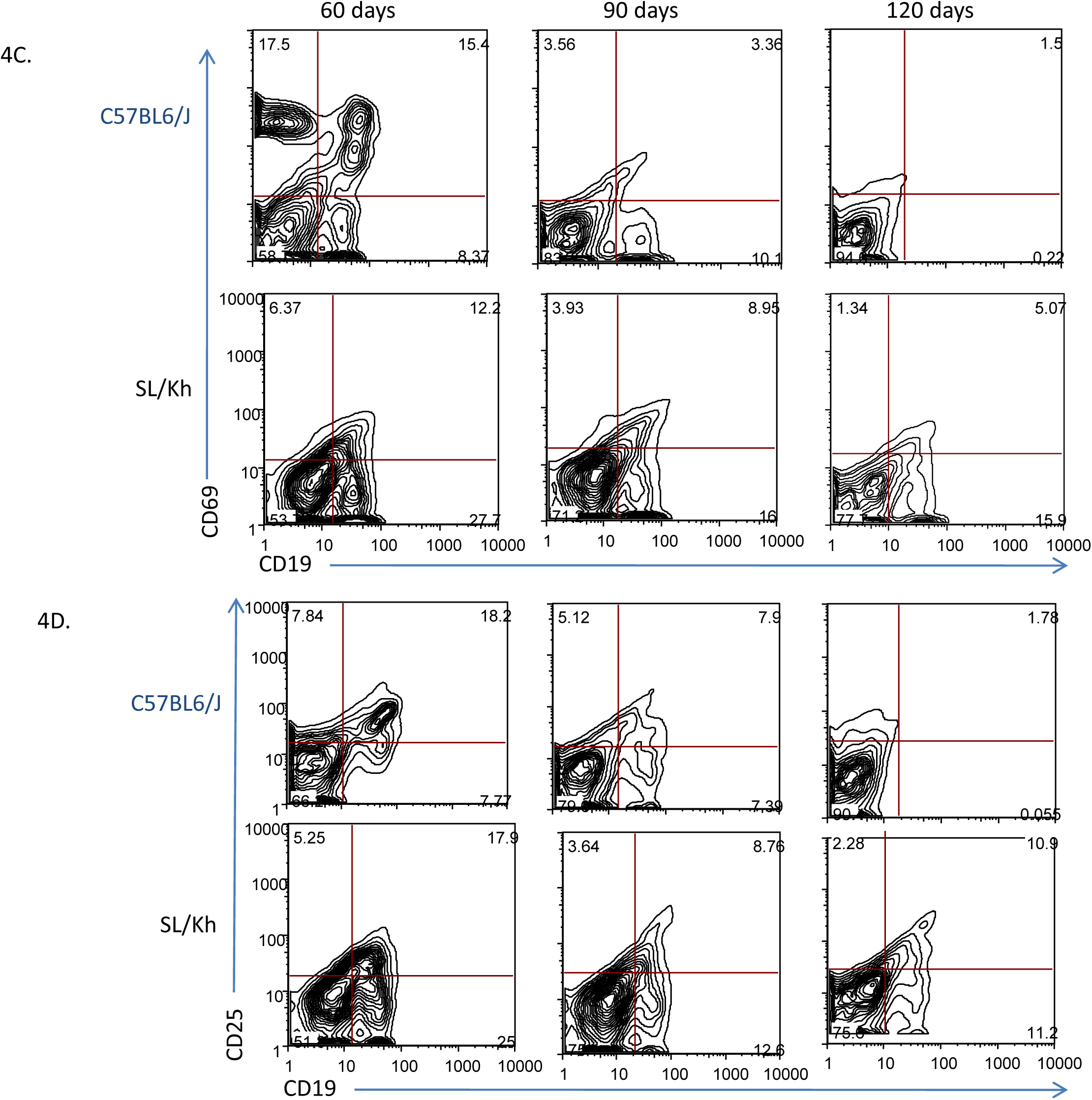
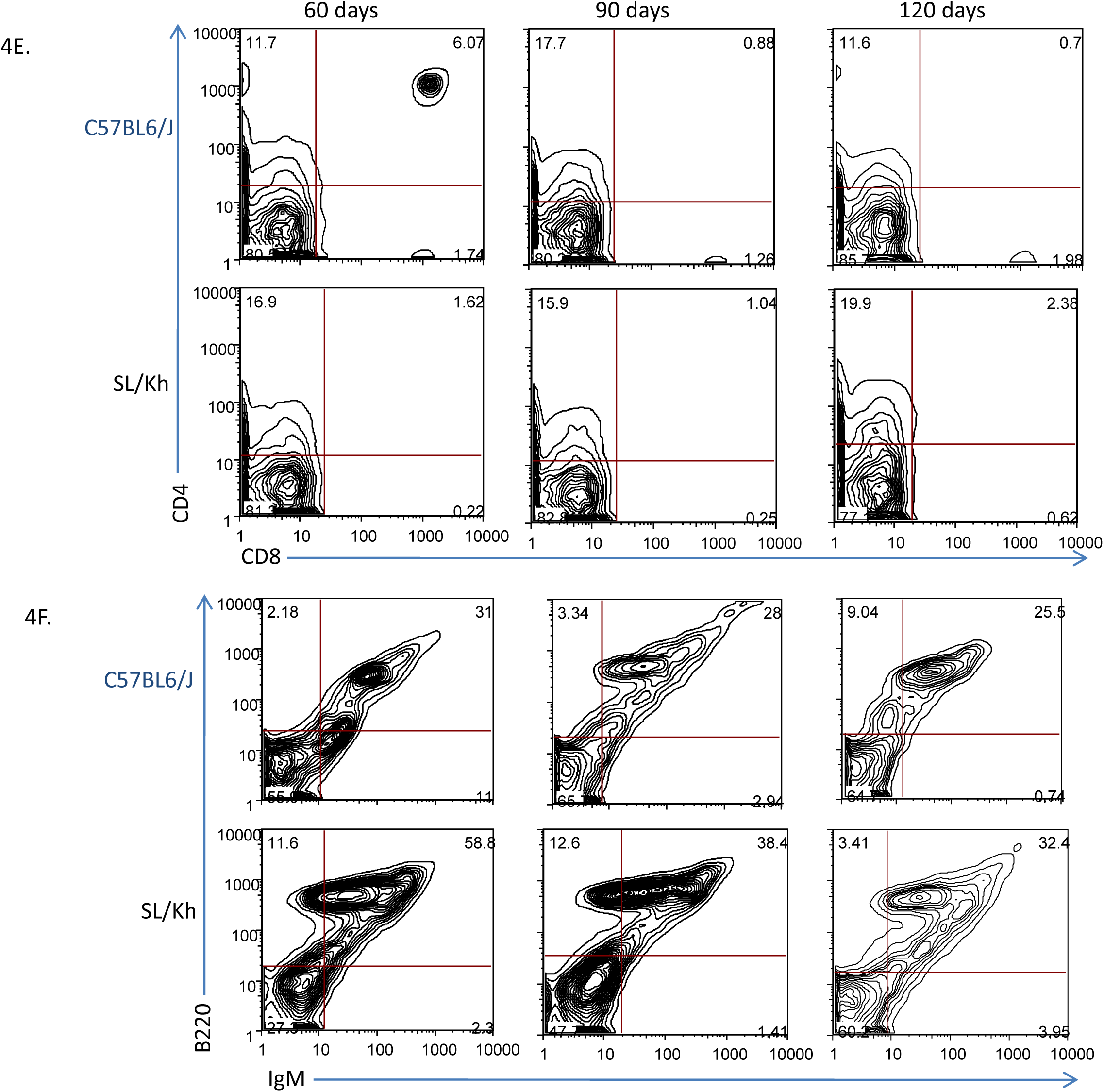
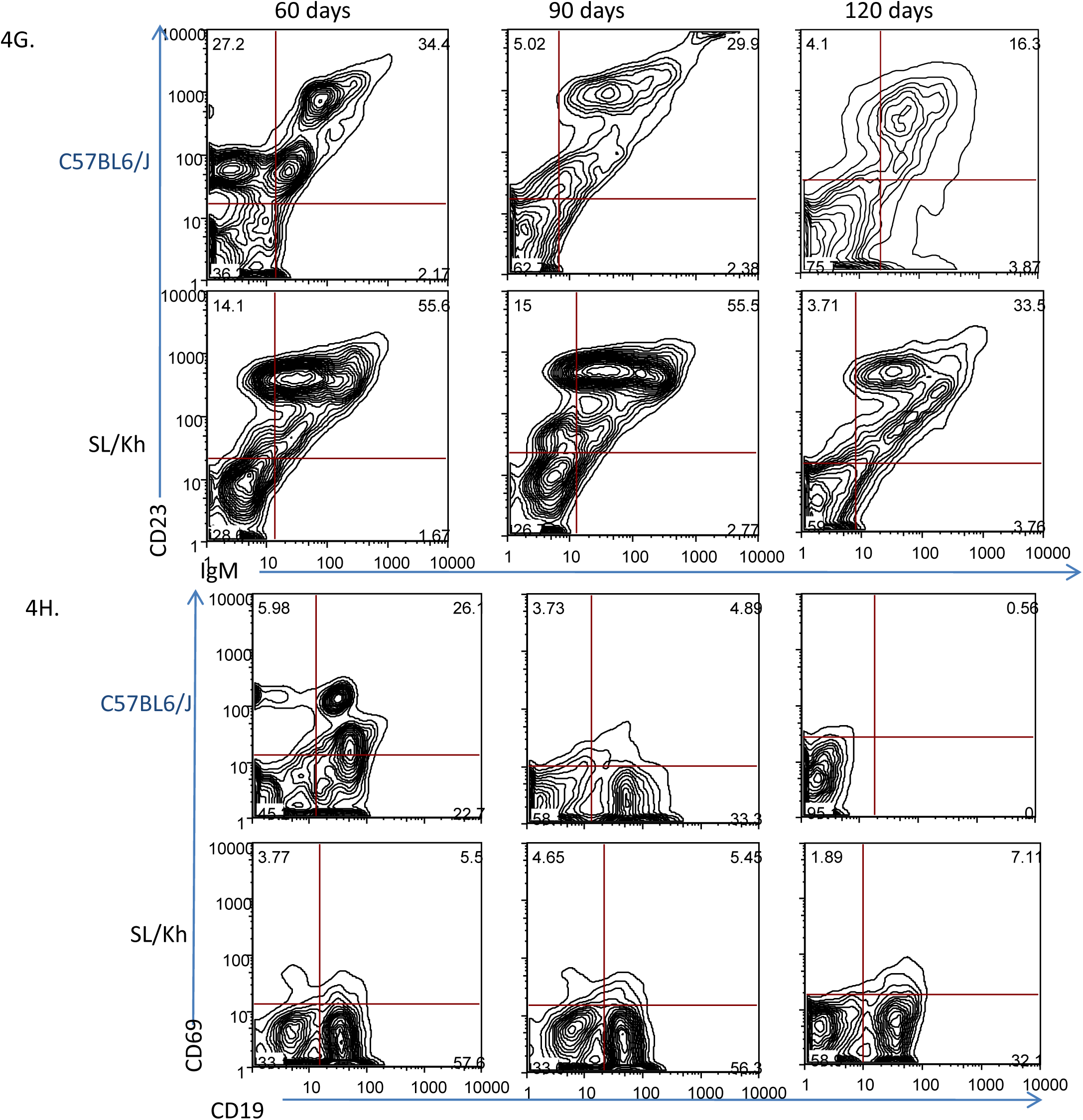
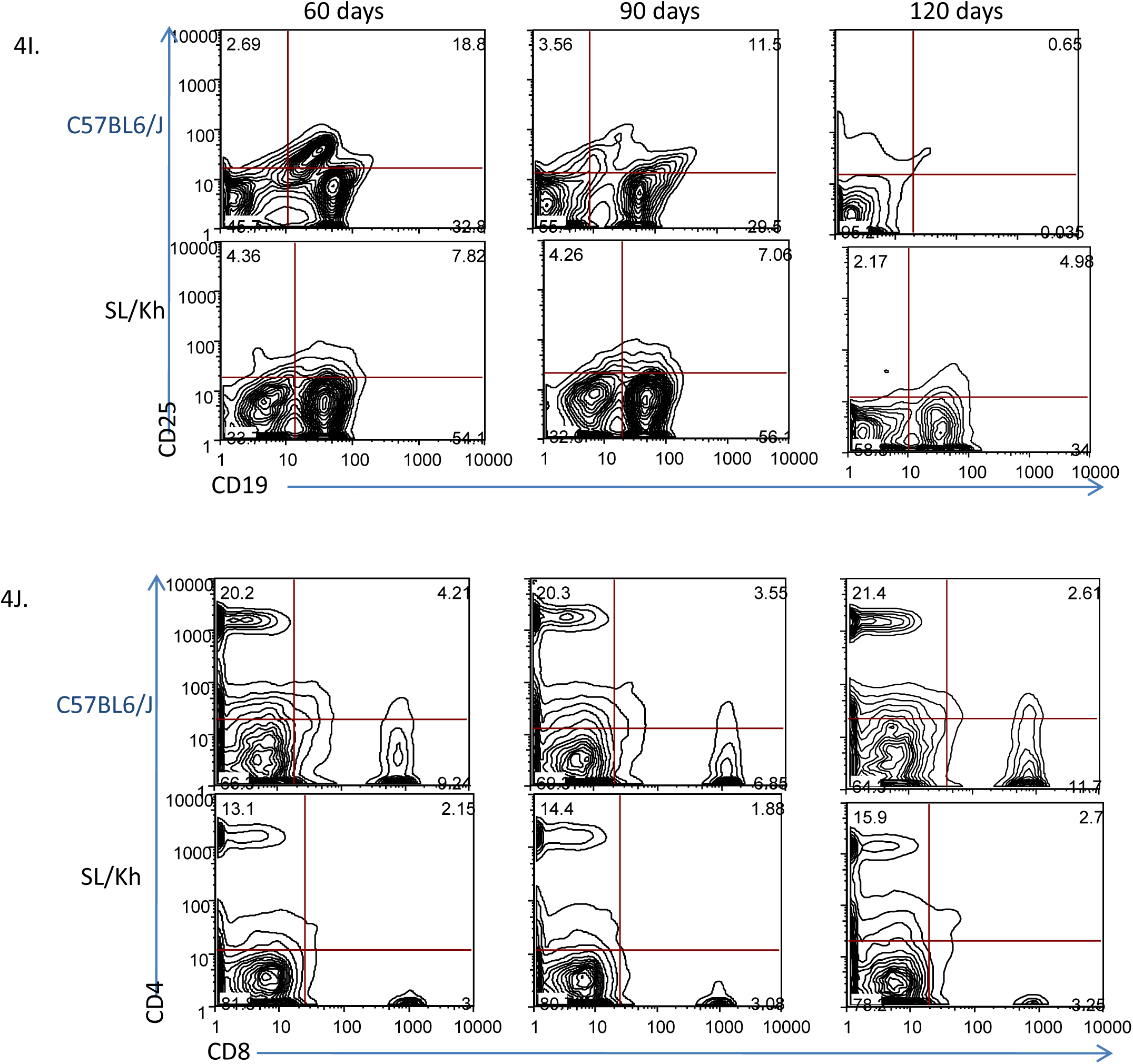
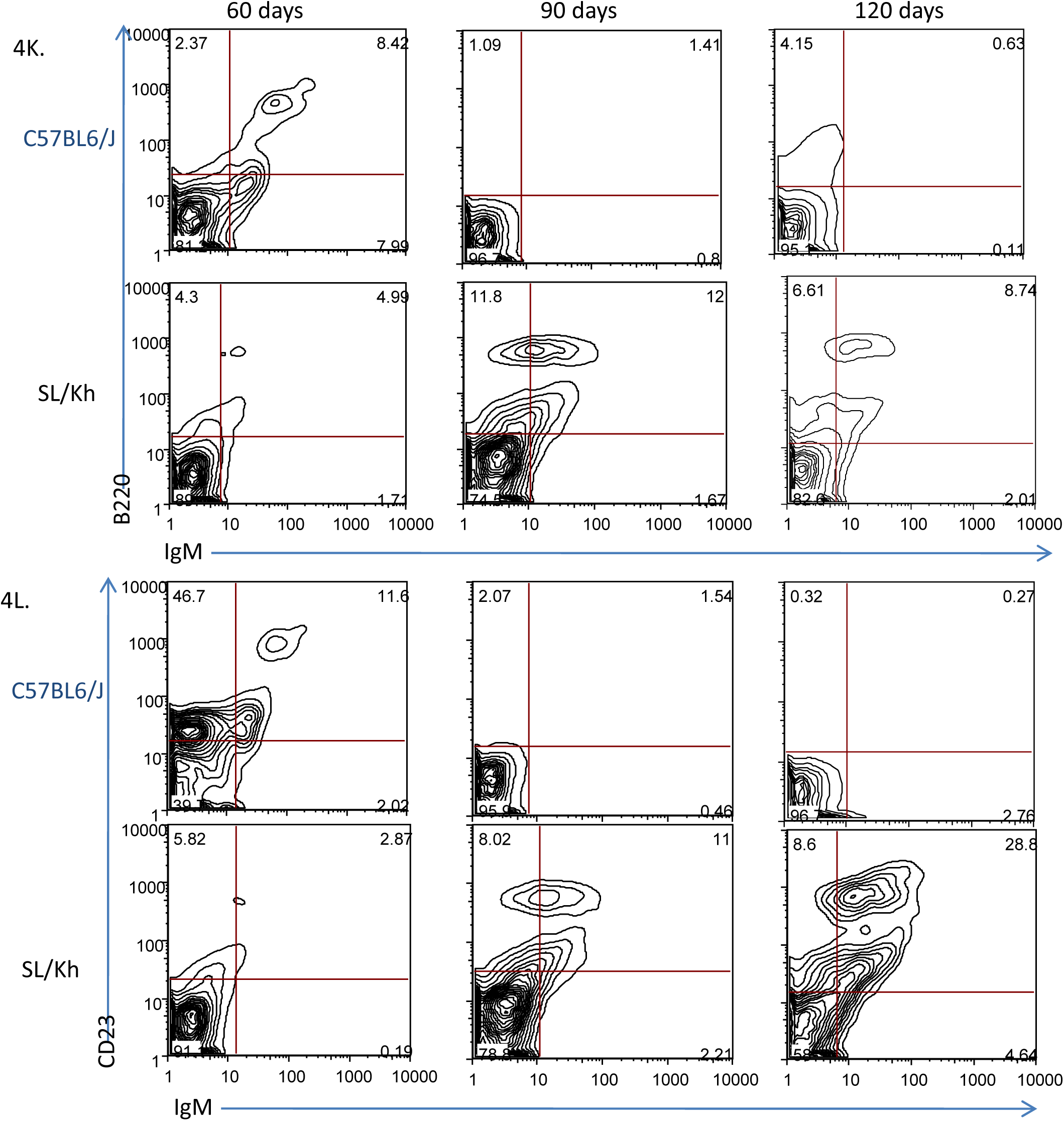
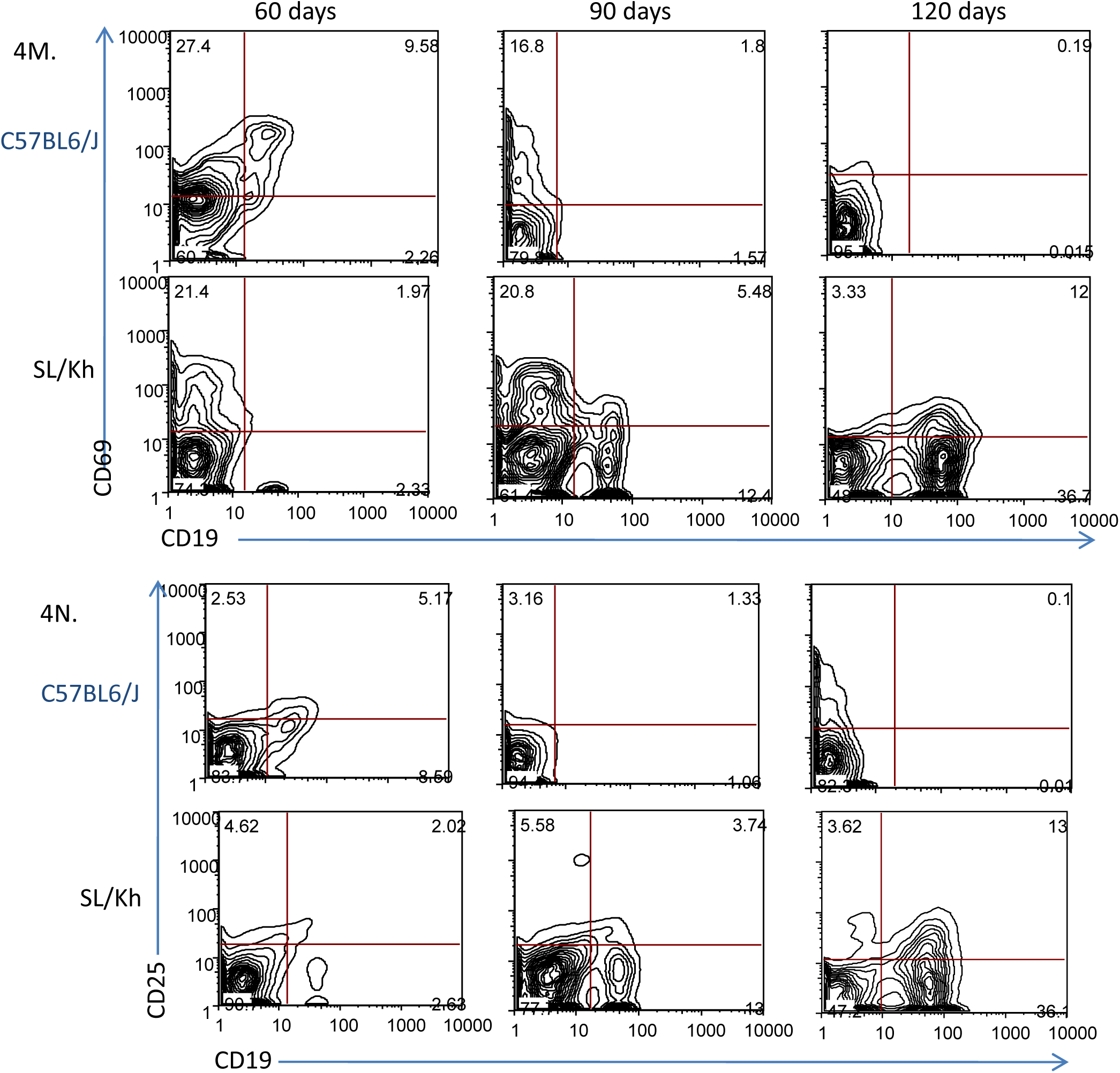
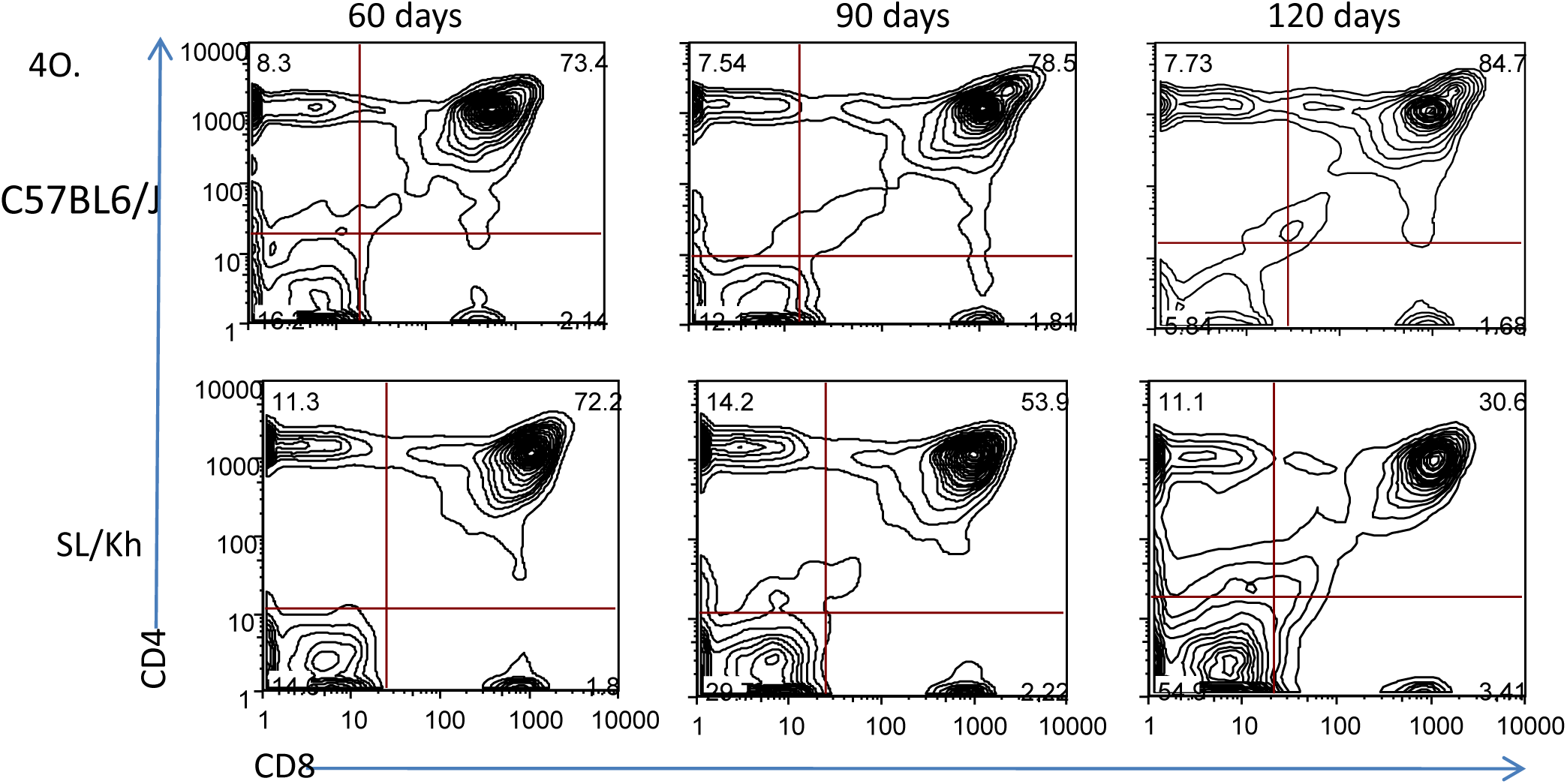
FACs analysis showing the lymphocyte landscape of SL/Kh and control C57BL6/J mice at 60, 90 and 120 days of age. Graphs are representative of the raw data collected and plotted in Figure 5. FACs data from bone marrow cells stained with B and T cell specific markers are shown in panels A-E, spleen cells in panels F-J and thymic cells in panels K-O. (A) Bone marrow cells stained with IgM and B220. (B) Bone marrow cells stained with B220 and CD23. (C) Bone marrow cells stained with CD19 and CD69. (D) Bone marrow cells stained with CD19 and CD25. (E) Bone marrow cells stained with CD8 and CD4. F. Spleen cells stained with IgM and B220. G. Spleen cells stained with IgM and CD23. H. Spleen cells stained with CD19 and CD69. I. Spleen cells stained with CD19 and CD25. J. Spleen cells stained with CD8 and CD4. K. Thymus cells stained with IgM and B220. L. Thymus cells stained with IgM and CD23. M. Thymus cells stained with CD19 and CD69. N. Thymus cells stained with CD19 and CD25. O. Thymus cells stained with CD8 and CD4.

---

**Supplementary Table 1.**
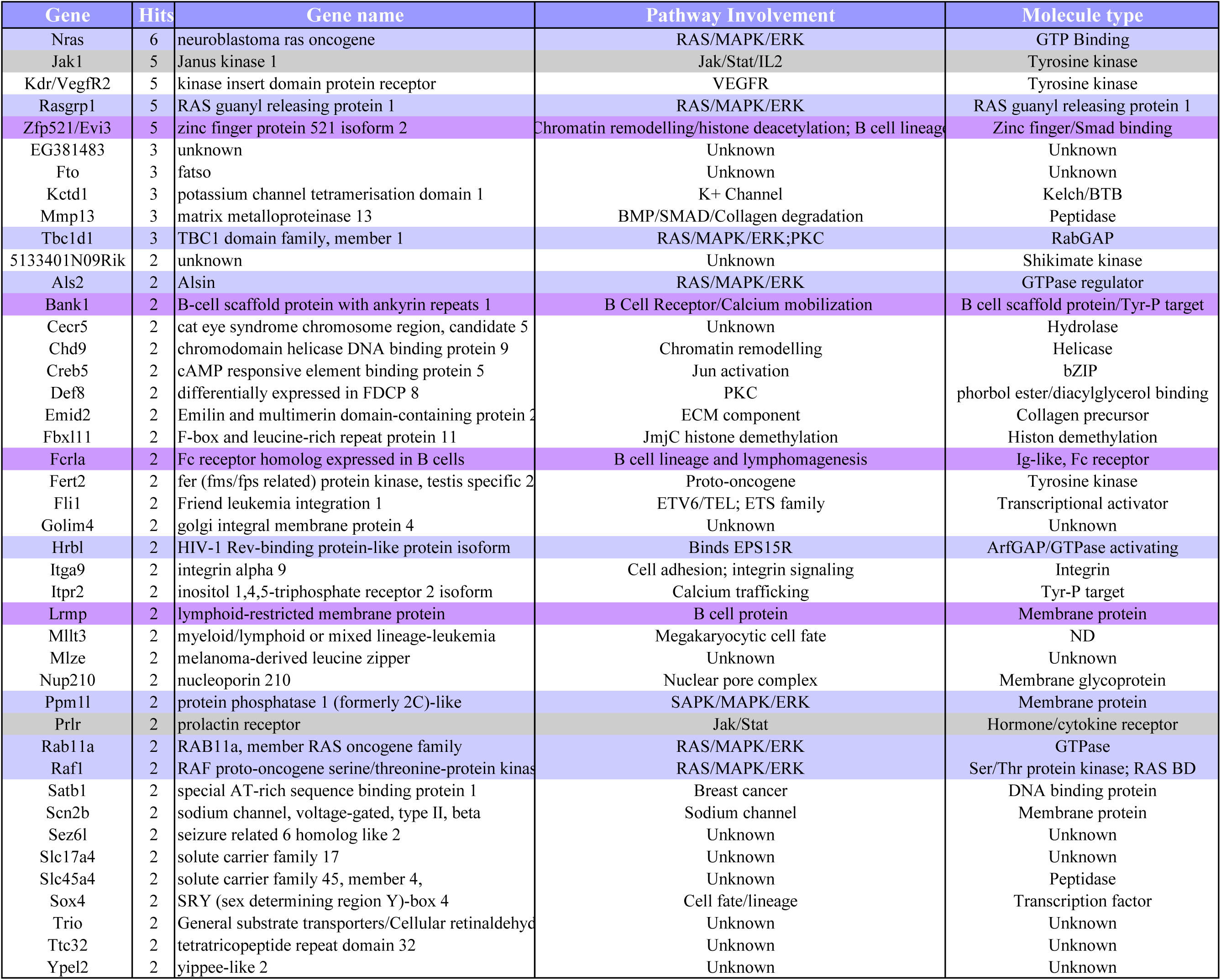
List of genes that are insertional mutated in two or more SL/Kh lymphomas. Many of the insertionally mutated genes function in the RAS/MAPK/ERK and JAK/STAT cell signaling pathways. The genes that function in the RAS/MAPK/ERK pathway are highlighted in light purple, the genes that function in the JAK/STAT pathway are highlighted in grey and the genes that are thought to play a role in the B cell pathway are highlighted in dark purple.

## References

1. Macdowell EC: Mouse leukemia. XVI. Spontaneous cases in strain C58 resisted by milk of old STOLI foster nurses, Cancer Res 1955, 15:23–25

2. Cole RKAF, J.: Experimental studies on the genetics of spontaneous leukemia in mice., Cancer Res 1941, 1:957–965

3. Anderson GW, Plagemann PG: Expression of ecotropic murine leukemia virus in the brains of C58/M, DBA2/J, and in utero-infected CE/J mice, J Virol 1995, 69:8089–8095

4. Contag CH, Plagemann PG: Susceptibility of C58 mice to paralytic disease induced by lactate dehydrogenase-elevating virus correlates with increased expression of endogenous retrovirus in motor neurons, Microb Pathog 1988, 5:287–296

5. Thomas CY, Roberts JS, Buxton VK: Mechanism of selection of class II recombinant murine leukemia viruses in the highly leukemic strain CWD, J Virol 1988, 62:1158–1166

6. Cloyd MW, Chattopadhyay SK: A new class of retrovirus present in many murine leukemia systems, Virology 1986, 151:31–40

7. Fredrickson TN, Morse HC, 3rd, Rowe WP: Spontaneous tumors of NFS mice congenic for ecotropic murine leukemia virus induction loci, J Natl Cancer Inst 1984, 73:521–524

8. Chieco-Bianchi L, Colombatti A, Collavo D, Sendo F, Aoki T, Fischinger PJ: Tumor induction by murine sarcoma virus in AKR and C58 mice. Reduction of tumor regression associated with appearance of Gross leukemia virus pseudotypes, J Exp Med 1974, 140:1162–1179

9. Zhang F, Schmidt WG, Hou Y, Williams AF, Jacobson K: Spontaneous incorporation of the glycosyl-phosphatidylinositol-linked protein Thy-1 into cell membranes, Proc Natl Acad Sci U S A 1992, 89:5231–5235

10. Hiai H, Kaneshima H, Nakamura H, Oguro YB, Moriwaki K, Nishizuka Y: Unusually early and high rate of spontaneous occurrence of nonthymic leukemias in SL/Kh mice, a subline of SL strain, Gann 1982, 73:704–712

11. Shimada MO, Yamada Y, Nakakuki Y, Okamoto K, Fukumoto M, Honjo T, Hiai H: SL/KH strain of mice: a model of spontaneous pre-B-lymphomas, Leuk Res 1993, 17:573–578

12. Okamoto K, Yamada Y, Ogawa MS, Toyokuni S, Nakakuki Y, Ikeda H, Yoshida O, Hiai H: Abnormal bone marrow B-cell differentiation in pre-B lymphoma-prone SL/Kh mice, Cancer Res 1994, 54:399–402

13. Hiai H, Tsuruyama T, Yamada Y: Pre-B lymphomas in SL/Kh mice: a multifactorial disease model, Cancer Sci 2003, 94:847–850

14. Yamada Y, Shisa H, Matsushiro H, Kamoto T, Kobayashi Y, Kawarai A, Hiai H: T lymphomagenesis is determined by a dominant host gene thymic lymphoma susceptible mouse-1 (TLSM-1) in mouse models, J Exp Med 1994, 180:2155–2162

15. Hiai H, Yamada Y, Shisa H, Kamoto T, Abujiang P, Lu LM: To be or not to be a T-lymphomas, that is determined by a dominant gene Tlsm-1 in mouse models, Leukemia 1997, 11 Suppl 3:193–194

16. Langdon WY, Harris AW, Cory S, Adams JM: The c-myc oncogene perturbs B lymphocyte development in E-mu-myc transgenic mice, Cell 1986, 47:11–18

17. Tidmarsh GF, Dailey MO, Whitlock CA, Pillemer E, Weissman IL: Transformed lymphocytes from Abelson-diseased mice express levels of a B lineage transformation-associated antigen elevated from that found on normal lymphocytes, J Exp Med 1985, 162:1421–1434

18. Hiai H, Buma YO, Ikeda H, Moriwaki K, Nishizuka Y: Epigenetic control of endogenous ecotropic virus expression in SL/Ni mice, J Natl Cancer Inst 1987, 79:781–787

19. Lu LM, Shimada R, Higashi S, Zeng Z, Hiai H: Bone marrow pre-B-1 (Bomb1): a quantitative trait locus inducing bone marrow pre-B-cell expansion in lymphoma-prone SL/Kh mice, Cancer Res 1999, 59:2593–2595

20. Du Y, Spence SE, Jenkins NA, Copeland NG: Cooperating cancer-gene identification through oncogenic-retrovirus-induced insertional mutagenesis, Blood 2005, 106:2498–2505

21. Godfrey DI, Kennedy J, Suda T, Zlotnik A: A developmental pathway involving four phenotypically and functionally distinct subsets of CD3-CD4-CD8-triple-negative adult mouse thymocytes defined by CD44 and CD25 expression, J Immunol 1993, 150:4244–4252

22. Hardy RR, Carmack CE, Shinton SA, Kemp JD, Hayakawa K: Resolution and characterization of pro-B and pre-pro-B cell stages in normal mouse bone marrow, J Exp Med 1991, 173:1213–1225

23. Laky K, Fowlkes BJ: Notch signaling in CD4 and CD8 T cell development, Curr Opin Immunol 2008, 20:197–202

24. Radtke F, Wilson A, MacDonald HR: Notch signaling in T- and B-cell development, Curr Opin Immunol 2004, 16:174–179

25. Deftos ML, Bevan MJ: Notch signaling in T cell development, Curr Opin Immunol 2000, 12:166–172

26. Wilson A, MacDonald HR, Radtke F: Notch 1-deficient common lymphoid precursors adopt a B cell fate in the thymus, J Exp Med 2001, 194:1003–1012

27. Erhardt I, Lischke A, Sebald W, Friedrich K: Constitutive association of JAK1 and STAT5 in pro-B cells is dissolved by interleukin-4-induced tyrosine phosphorylation of both proteins, FEBS Lett 1998, 439:71–74

28. Iritani BM, Forbush KA, Farrar MA, Perlmutter RM: Control of B cell development by Ras-mediated activation of Raf, EMBO J 1997, 16:7019–7031

29. Mangues R, Symmans WF, Lu S, Schwartz S, Pellicer A: Activated N-ras oncogene and N-ras proto-oncogene act through the same pathway for in vivo tumorigenesis, Oncogene 1996, 13:1053–1063

30. Verma A, Kambhampati S, Parmar S, Platanias LC: Jak family of kinases in cancer, Cancer Metastasis Rev 2003, 22:423–434

31. Tokoro Y, Sugawara T, Yaginuma H, Nakauchi H, Terhorst C, Wang B, Takahama Y: A mouse carrying genetic defect in the choice between T and B lymphocytes, J Immunol 1998, 161:4591–4598

32. Radtke F, Wilson A, Stark G, Bauer M, van Meerwijk J, MacDonald HR, Aguet M: Deficient T cell fate specification in mice with an induced inactivation of Notch1, Immunity 1999, 10:547–558

33. Akashi K, Richie LI, Miyamoto T, Carr WH, Weissman IL: B lymphopoiesis in the thymus, J Immunol 2000, 164:5221–5226

34. Campos-Ortega JA: Mechanisms of early neurogenesis in Drosophila melanogaster, J Neurobiol 1993, 24:1305–1327

35. Doe CQ, Goodman CS: Early events in insect neurogenesis. II. The role of cell interactions and cell lineage in the determination of neuronal precursor cells, Dev Biol 1985, 111:206–219

36. Turcotte K, Gauthier S, Tuite A, Mullick A, Malo D, Gros P: A mutation in the Icsbp1 gene causes susceptibility to infection and a chronic myeloid leukemia-like syndrome in BXH-2 mice, J Exp Med 2005, 201:881–890

37. Turcotte K, Gauthier S, Malo D, Tam M, Stevenson MM, Gros P: Icsbp1/IRF-8 is required for innate and adaptive immune responses against intracellular pathogens, J Immunol 2007, 179:2467–2476

38. Wu L, Antica M, Johnson GR, Scollay R, Shortman K: Developmental potential of the earliest precursor cells from the adult mouse thymus, J Exp Med 1991, 174:1617–1627

39. Matsuzaki Y, Gyotoku J, Ogawa M, Nishikawa S, Katsura Y, Gachelin G, Nakauchi H: Characterization of c-kit positive intrathymic stem cells that are restricted to lymphoid differentiation, J Exp Med 1993, 178:1283–1292

40. Wu L, Li CL, Shortman K: Thymic dendritic cell precursors: relationship to the T lymphocyte lineage and phenotype of the dendritic cell progeny, J Exp Med 1996, 184:903–911

41. Weissman IL: Thymus cell maturation. Studies on the origin of cortisone-resistant thymic lymphocytes, J Exp Med 1973, 137:504–510

42. Lu LM, Hiai H: Mixed phenotype lymphomas in thymectomized (SL/KhxAKR/Ms)F1 mice, Jpn J Cancer Res 1999, 90:1218–1223

43. Levitt LJ, Aisenberg AC, Harris NL, Linggood RM, Poppema S: Primary non-Hodgkin’s lymphoma of the mediastinum, Cancer 1982, 50:2486–2492

44. Trump DL, Mann RB: Diffuse large cell and undifferentiated lymphomas with prominent mediastinal involvement, Cancer 1982, 50:277–282

45. Harris NL, Jaffe ES, Stein H, Banks PM, Chan JK, Cleary ML, Delsol G, De Wolf-Peeters C, Falini B, Gatter KC, et al.: A revised European-American classification of lymphoid neoplasms: a proposal from the International Lymphoma Study Group, Blood 1994, 84:1361–1392

46. al-Sharabati M, Chittal S, Duga-Neulat I, Laurent G, Mazerolles C, al-Saati T, Brousset P, Delsol G: Primary anterior mediastinal B-cell lymphoma. A clinicopathologic and immunohistochemical study of 16 cases, Cancer 1991, 67:2579–2587

47. Hofmann WJ, Momburg F, Moller P: Thymic medullary cells expressing B lymphocyte antigens, Hum Pathol 1988, 19:1280–1287

48. Isaacson PG, Norton AJ, Addis BJ: The human thymus contains a novel population of B lymphocytes, Lancet 1987, 2:1488–1491

49. Kanavaros P, Gaulard P, Charlotte F, Martin N, Ducos C, Lebezu M, Mason DY: Discordant expression of immunoglobulin and its associated molecule mb-1/CD79a is frequently found in mediastinal large B cell lymphomas, Am J Pathol 1995, 146:735–741

50. Benschop RJ, Cambier JC: B cell development: signal transduction by antigen receptors and their surrogates, Curr Opin Immunol 1999, 11:143–151

51. Peschon JJ, Morrissey PJ, Grabstein KH, Ramsdell FJ, Maraskovsky E, Gliniak BC, Park LS, Ziegler SF, Williams DE, Ware CB, Meyer JD, Davison BL: Early lymphocyte expansion is severely impaired in interleukin 7 receptor-deficient mice, J Exp Med 1994, 180:1955–1960

52. Cao X, Shores EW, Hu-Li J, Anver MR, Kelsall BL, Russell SM, Drago J, Noguchi M, Grinberg A, Bloom ET, et al.: Defective lymphoid development in mice lacking expression of the common cytokine receptor gamma chain, Immunity 1995, 2:223–238

53. Nosaka T, van Deursen JM, Tripp RA, Thierfelder WE, Witthuhn BA, McMickle AP, Doherty PC, Grosveld GC, Ihle JN: Defective lymphoid development in mice lacking Jak3, Science 1995, 270:800–802

54. Rosenwald A, Wright G, Leroy K, Yu X, Gaulard P, Gascoyne RD, Chan WC, Zhao T, Haioun C, Greiner TC, Weisenburger DD, Lynch JC, Vose J, Armitage JO, Smeland EB, Kvaloy S, Holte H, Delabie J, Campo E, Montserrat E, Lopez-Guillermo A, Ott G, Muller-Hermelink HK, Connors JM, Braziel R, Grogan TM, Fisher RI, Miller TP, LeBlanc M, Chiorazzi M, Zhao H, Yang L, Powell J, Wilson WH, Jaffe ES, Simon R, Klausner RD, Staudt LM: Molecular diagnosis of primary mediastinal B cell lymphoma identifies a clinically favorable subgroup of diffuse large B cell lymphoma related to Hodgkin lymphoma, J Exp Med 2003, 198:851–862

55. Savage KJ, Monti S, Kutok JL, Cattoretti G, Neuberg D, De Leval L, Kurtin P, Dal Cin P, Ladd C, Feuerhake F, Aguiar RC, Li S, Salles G, Berger F, Jing W, Pinkus GS, Habermann T, Dalla-Favera R, Harris NL, Aster JC, Golub TR, Shipp MA: The molecular signature of mediastinal large B-cell lymphoma differs from that of other diffuse large B-cell lymphomas and shares features with classical Hodgkin lymphoma, Blood 2003, 102:3871–3879

56. Melzner I, Bucur AJ, Bruderlein S, Dorsch K, Hasel C, Barth TF, Leithauser F, Moller P: Biallelic mutation of SOCS-1 impairs JAK2 degradation and sustains phospho-JAK2 action in the MedB-1 mediastinal lymphoma line, Blood 2005, 105:2535–2542

57. Cacalano NA, Sanden D, Johnston JA: Tyrosine-phosphorylated SOCS-3 inhibits STAT activation but binds to p120 RasGAP and activates Ras, Nat Cell Biol 2001, 3:460–465

58. Krauth J: The interpretation of significance tests for independent and dependent samples, J Neurosci Methods 1983, 9:269–281

59. Tsuruyama T, Nakamura T, Jin G, Ozeki M, Yamada Y, Hiai H: Constitutive activation of Stat5a by retrovirus integration in early pre-B lymphomas of SL/Kh strain mice, Proc Natl Acad Sci U S A 2002, 99:8253–8258

60. Jin G, Tsuruyama T, Yamada Y, Hiai H: Svi3: A provirus common integration site in c-myc in SL/Kh pre-B lymphomas, Cancer Sci 2003, 94:791–795

